# Viral infection engenders bona fide and bystander lung memory B cell subsets through permissive selection

**DOI:** 10.1101/2021.12.14.472614

**Authors:** Claude Gregoire, Lionel Spinelli, Sergio Villazala-Merino, Laurine Gil, Myriam Moussa, Chuang Dong, Ana Zarubica, Mathieu Fallet, Jean-Marc Navarro, Bernard Malissen, Pierre Milpied, Mauro Gaya

**Author notes:** Correspondance (P.M.), (M.G.).

## Abstract

Lung-resident memory B cells (MBCs) provide localized protection against reinfection in the respiratory airways. Currently, the biology of these cells remains largely unexplored. Here, we combined influenza and SARS-CoV-2 infection with fluorescent-reporter mice to identify MBCs regardless of antigen specificity. scRNA-seq analysis and confocal imaging revealed that two main transcriptionally distinct subsets of MBCs colonize the lung peribronchial niche after infection. These subsets arise from different progenitors and are both class-switched, somatically mutated and intrinsically biased in their differentiation fate towards plasma cells. Combined analysis of antigen-specificity and B cell receptor repertoire unveiled a highly permissive selection process that segregates these subsets into *“bona fide”* virus-specific MBCs and *“bystander”* MBCs with no apparent specificity for eliciting viruses. Thus, diverse transcriptional programs in MBCs are not linked to specific effector fates but rather to divergent strategies of the immune system to simultaneously provide rapid protection from reinfection while diversifying the initial B cell repertoire.

## Introduction

The immune system mounts protective responses to counteract the threat posed by pathogens during infection. These responses leave immunological memory, a strategy that allows our body to remember previously encountered pathogens. Memory B cells (MBCs) and T cells are long-lived lymphocytes that constitute an essential component of this strategy (Akkaya et al., 2020). These cells take up residence in secondary lymphoid organs and remain in quiescent state until a secondary antigen encounter. Upon re-challenge, they rapidly produce large numbers of effector cells that deliver fast and effective protection (Weisel and Shlomchik, 2017). In the case of MBCs, they can either differentiate into short-lived plasma cells, which produce high-affinity neutralizing antibodies, or re-enter germinal centers and provide, up to a certain extent, a new source of long-lasting protection with increased affinity and breadth (Kurosaki et al., 2015; McHeyzer-Williams et al., 2015; Mesin et al., 2020; Viant et al., 2020). MBC fate decision can be shaped by cell-intrinsic features, B cell receptor isotype, antigen affinity and the magnitude of CD40 signalling (Dogan et al., 2009; Koike et al., 2019; Pape et al., 2011; Viant et al., 2020, 2021; Zuccarino-Catania et al., 2014).

Memory lymphocytes do not exclusively reside in secondary lymphoid organs. For instance, a lineage of T cells can occupy the tissue barriers after infection without recirculating. These tissue-resident memory T cells are transcriptionally and functionally distinct from recirculating ones, and provide site-specific responses against infection (Szabo et al., 2019). Yet, if a B cell counterpart of these cells existed remained unknown. A recent study has shown, through the use of elegant parabiotic experiments, that a population of tissue-resident MBCs settle in the lungs after influenza virus infection (Allie et al., 2019). Lung MBCs not only ensure a first layer of protection directly at the tissue barrier but also display high cross-reactivity to viral escapes, highlighting the potential of targeting them to develop broadly protective vaccines (Adachi et al., 2015; Onodera et al., 2012). At present, it remains unknown if lung MBCs occupy specific tissue niches in the lung mucosa, if they bear special transcriptional programs that allow their survival in the lung airways or if discrete MBC subsets coexist upon infection.

Here, we combined the use of Aicda-Cre^ERT2^ Rosa26-EYFP reporter mice with influenza and SARS-CoV-2 infection models to track lung MBCs based on past Aicda expression at the time of viral infection. As this approach does not rely on the ability of MBCs to bind viral antigens, it gave us access to cells with little or no affinity for specific antigens, which can represent a large proportion of MBCs (Viant et al., 2020) and were disregarded in previous studies. We show, for the first time, that two main subsets of MBCs permanently colonize the lung peribronchial niche upon respiratory viral infection. These subsets arise from different progenitors that underwent class-switching and somatic hypermutation in germinal centers, and subsequently acquired divergent transcriptional programs and chemokine/Fc receptor patterns. Unexpectedly, both MBC subsets are intrinsically biased in their differentiation fate towards plasma cells, highlighting that divergent transcriptional programs are not associated with specific effector fates. Instead, these subsets segregate “*bona fide*” MBCs, which dominate recall responses by producing high affinity antibody-secreting cells, from “*bystander*” MBCs, with no specificity for the immunogen, unable to produce protective antibodies but with the ability to retain and display antigen in the form of immune complexes. These results challenge the notion that germinal centers only work as “machines” to produce high-affinity *bona fide* MBCs. Instead, permissive selection mechanisms simultaneously operate in these reactions, expanding the diversity of the initial B cell repertoire and giving rise to *bystander* MBCs.

## Results

### Unbiased identification of MBCs homing to the lungs and lymphoid organs upon influenza infection

In order to identify lung-homing MBCs without introducing a bias on antigen specificity, we took advantage of the Aicda-Cre^ERT2^ Rosa26-EYFP mouse strain, from now on referred as Aid-EYFP (Le Gallou et al., 2018). This model allows us to permanently label cells that have expressed activation-induced cytidine deaminase, the enzyme that initiates class-switching and somatic hypermutation, at the time of tamoxifen treatment. As a vast fraction of MBCs have gone through at least one of these two processes early in the response to infection, we expect them to be labelled by tamoxifen. Accordingly, we intranasally infected Aid-EYFP mice with 5 plaque-forming units (PFUs) of influenza A virus Puerto Rico/8/34 (PR8) H1N1 strain, and orally administered tamoxifen at days 6, 8 and 10. We assessed the efficiency and specificity of this strategy after 10 weeks of infection by flow cytometry (Figure 1A). We observed the presence of a discrete population of YFP^+^ cells in lungs and secondary lymphoid organs, mostly corresponding to CD19^+^ B cells and a small population of CD19^-^ plasma cells (Figure 1B and Extended Figure 1A). Strikingly, more than 90% of YFP^+^ lung cells showed a phenotype associated with MBCs (CD38^+^ GL-7^-^) (Figure 1C). In contrast, only 30% of YFP^+^ lymph node cells showed a MBC phenotype; the great majority were still committed to long-lived germinal centers (CD38^-^ GL-7^+^) that persist in mediastinal lymph nodes long after viral clearance (Figure 1C).

**Figure 1.**
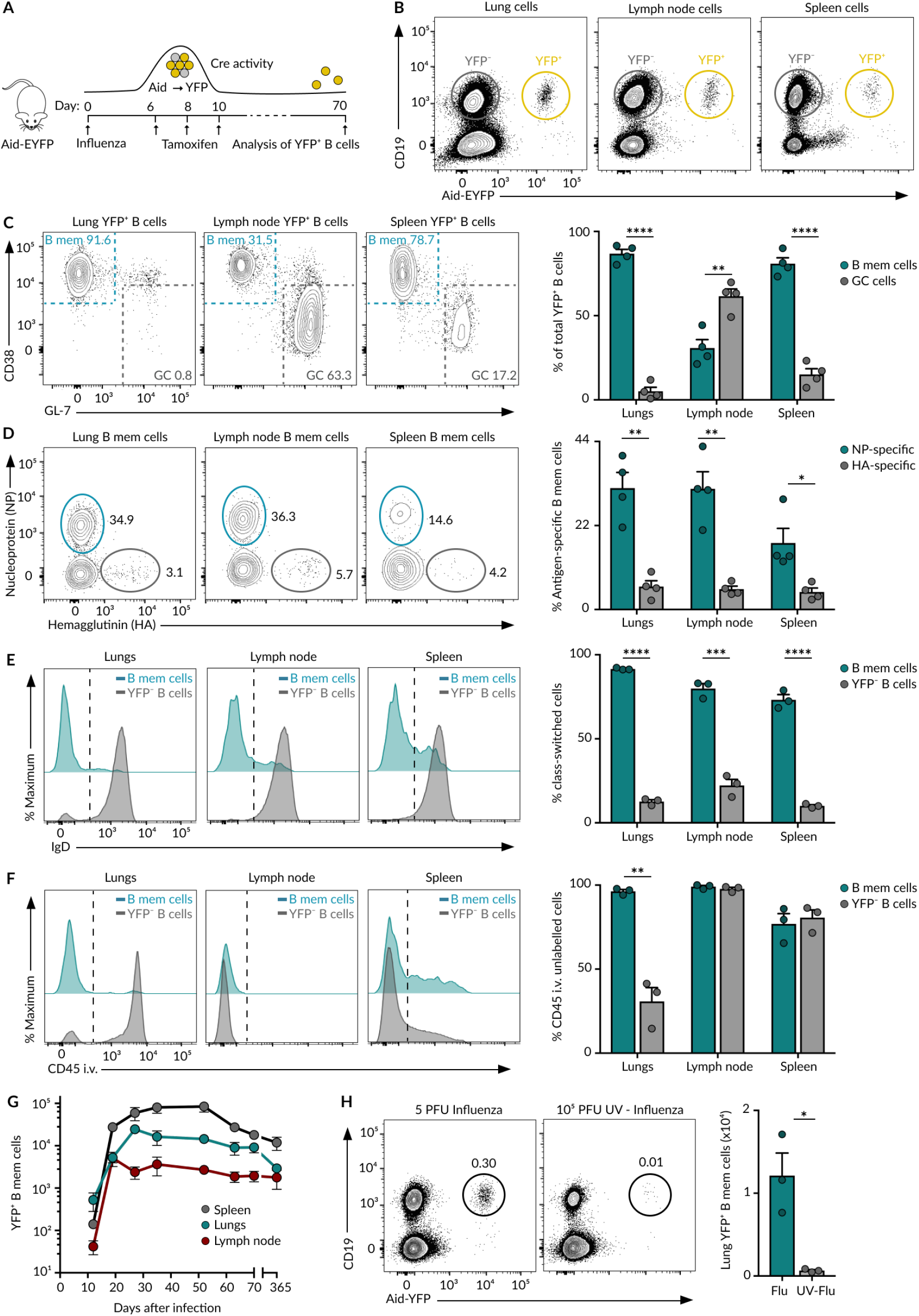
MBCs in influenza infection. **(A)** Overview of experimental approach. **(B)** Flow cytometry plots displaying YFP+ and YFP-B cells in lungs, mediastinal lymph nodes and spleen of Aid-EYFP animals treated as in A. **(C)** Flow cytometry plots showing the percentage of MBCs (CD38+GL7-) and germinal center (CD38-GL-7+) cells gated from the CD19+ YFP+ population in B. **(D)** Flow cytometry plots showing the percentage of YFP+ MBCs that bind to fluorescently labelled Nucleoprotein (NP) and Hemagglutinin (HA). **(E)** Histograms depicting IgD expression by YFP+ MBCs and YFP-B cells. **(F)** Histograms showing the extent of labelling of YFP+ MBCs and YFP-B cells with anti-CD45 antibody administered intravenously 5 minutes before sacrifice. Quantification shows the percentage of B cells that are protected from in vivo staining. **(G)** Absolute number of YFP+ MBCs in lungs, lymph nodes and spleen of Aid-EYFP mice treated as in A and analyzed at indicated time points. **(H)** Flow cytometry plots displaying YFP+ B cells in lungs of Aid-EYFP animals (day 70) infected with 5 PFU of influenza virus or challenged with 10^5^ PFU of influenza virus previously inactivated by UV light. In all panels, bar charts show the quantification of one representative experiment out of three, mean ± s.e.m. Each dot represents one mouse. t test: *p<0.05, **p<0.01, ***p<0.001 and ****p<0.0001.

To determine whether these YFP^+^ cells are genuine MBCs, we measured antigen-specificity, class-switching, accessibility to blood circulation and lifespan. Regarding specificity, we detected an average of 30% of YFP^+^ B cells binding to APC-labelled influenza nucleoprotein (NP) and an average of 5% that bound to PE-labelled influenza hemagglutinin (HA); less than 2% of YFP^-^ B cells bound to these proteins (Figure 1D and Extended Figure 1B). These results indicate that the YFP^+^ population is enriched in B cells specific for influenza antigens. Regarding class-switching, we observed that YFP^+^ cells lacked IgD expression while YFP^-^ cells were mainly IgD^+^, showing that YFP^+^ cells have undergone extensive class-switching (Figure 1E). To measure accessibility to blood, we administered fluorescently labelled anti-CD45 antibody intravenously 5 minutes before sacrifice. We found that more than 95% of lung YFP^+^ cells, in marked contrast to YFP^-^ B cells, were protected from *in vivo* antibody labelling, indicating that these cells preferentially accumulate in the lung parenchyma rather than in the blood circulation (Figure 1F). In mediastinal lymph nodes and spleen, both YFP^+^ and YFP^-^ were mostly protected from in vivo antibody labelling, in line with the notion that B cells residing in lymphoid organs are not in direct contact with the blood circulation (Figure 1F). As expected, only spleen marginal zone B cells were labelled by anti-CD45 (Figures 1F and Extended Figure 1C). We finally enumerated YFP^+^ cells in lungs, mediastinal lymph node and spleen at different time points after infection to get an insight into the lifespan of these cells. We found that the kinetics of YFP^+^ cells follow a similar trend across organs: YFP^+^ cells were detected at low numbers at day 12, peaked between days 20-30 and slowly declined thereafter (Figure 1G). Remarkably, YFP^+^ B cells were detected in significant numbers even one year after infection, suggesting that these cells are long-lived (Figure 1G). The accumulation of YFP^+^ B cells in lungs was not observed in mice administered with UV-inactivated influenza virus, pointing out that infection, or the inflammation process associated with it, is required for the establishment of MBCs (Figure 1H). Altogether, these results highlight the power of our strategy to track long-lived class-switched MBCs homing to the lungs and lymphoid organs without introducing a bias on antigen specificity.

### Transcriptionally distinct subsets of MBCs co-exist in lungs and lymphoid organs

To unveil if influenza infection engenders discrete subsets of MBCs in lungs and draining lymphoid organs, we performed scRNA-seq on YFP^+^CD19^+^CD38^+^GL-7^-^ MBCs sorted from lungs, mediastinal lymph nodes and spleen at day 70 after influenza infection. After tagging cells with mouse- and organ-specific hashtag antibodies, we captured single cells from three mice using the droplet-based microfluidic system Chromium (10X) and performed 5’-end scRNA-seq and scBCR-seq. After processing and filtering (see Methods), we retained 4,679 good quality cells for subsequent analysis: lung n=2,041; lymph node n=521; spleen n=2,117. We performed Uniform Manifold Approximation and Projection (UMAP) for dimensionality reduction within each tissue and found that MBCs bifurcated into three separate clusters within lungs and mediastinal lymph nodes and into five clusters in spleen (Figures 2A-2D). By analyzing gene expression, we identified groups of marker genes associated with specific B cell clusters, reflecting divergent transcriptional programs within these memory populations (Figures 2E). To compare the extent of similarity across lung MBC clusters with those from draining lymphoid organs, we calculated the Szymkiewicz Simpson similarity index. We found that lung clusters Lg1, Lg2 and Lg3 shared high levels of marker genes’ overlap with lymph node clusters Ln1, Ln2 and Ln3 respectively, indicating shared memory mechanisms operating across these organs (Figure 2F). In contrast, the spleen contained specific MBC populations, such as Sp4 and Sp5, that were absent in lungs (Figure 2F). These data show that the MBC pool is not homogeneous but, instead, is constituted of transcriptionally distinct subpopulations that are either tissue-specific or shared across organs.

**Figure 2.**
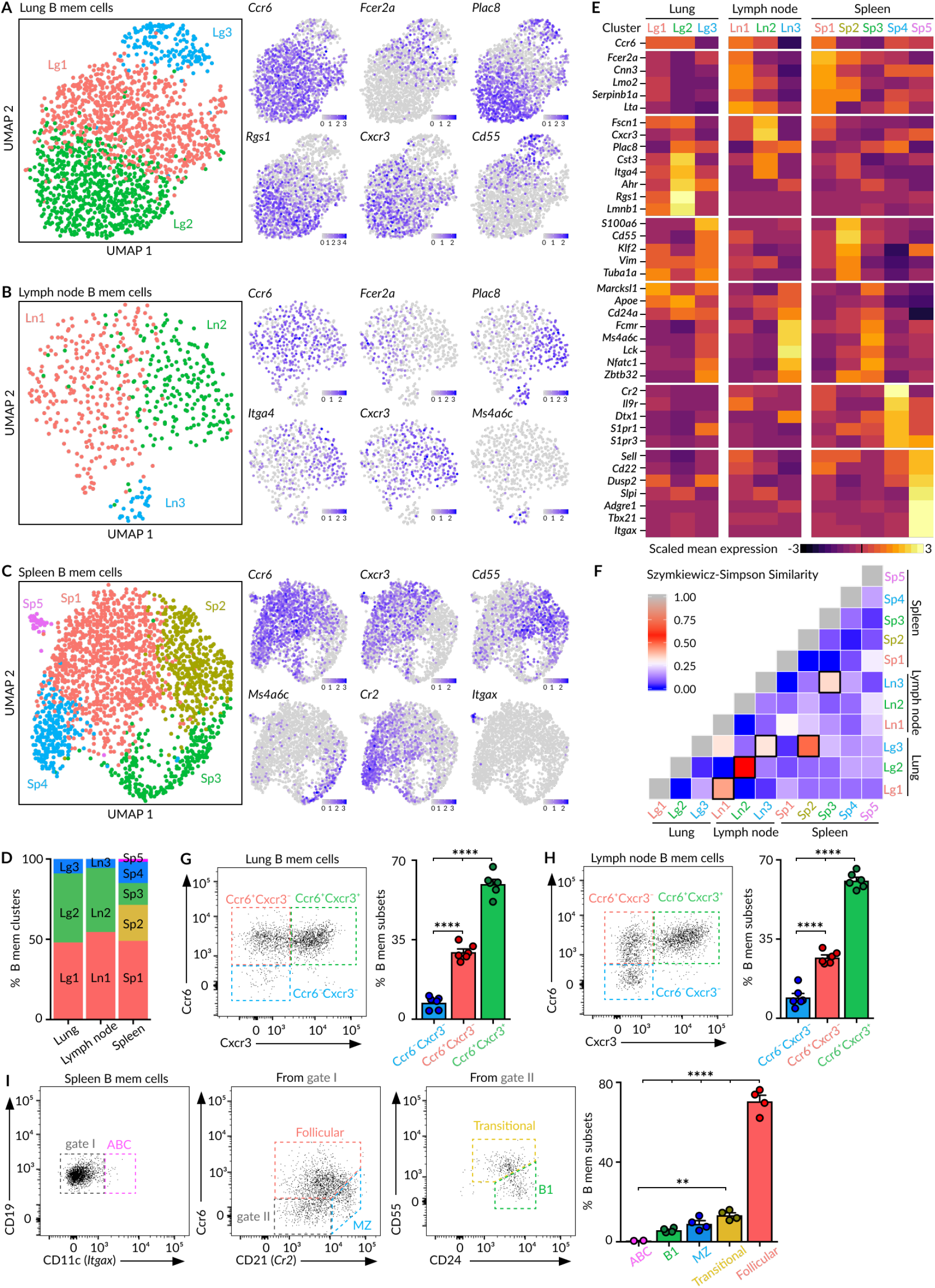
Heterogeneity of MBCs. **(A-C)** UMAP projections of MBCs in lungs (A), mediastinal lymph node (B) and spleen (C), colored by subpopulation. Feature plots display the expression of indicated marker genes in MBCs laid out in the UMAP representation. Scale: normalized UMI counts. **(D)** Bar chart showing the relative proportions of MBC subpopulations within each tissue. **(E)** Heatmaps exhibiting marker gene expression for each tissue subpopulation. Colour: Scaled mean expression. **(F)** Szymkiewicz-Simpson similarity matrix for pairwise comparisons among subpopulations from different tissues. Black rectangles indicate values higher than 0.32. **(G-H)** Flow cytometry plots showing the presence of Ccr6^-^Cxcr3^-^, Ccr6^+^Cxcr3^-^ and Ccr6^+^Cxcr3^+^ MBC subsets in the CD19^+^ YFP^+^CD38^+^GL7^-^ population from lungs (G) and lymph nodes (H). **(I)** Flow cytometry plots showing the gating strategy for spleen MBC subsets according to the expression of Cd11c, Ccr6, Cd21, Cd24 and Cd55. In panels G to I, bar charts show the quantification of one representative experiment out of three, mean ± s.e.m. Each dot represents one mouse. One-way Anova test: **p<0.01 and ****p<0.0001.

Among MBCs residing in lungs and lymph nodes, those belonging to the two major clusters (Lg1-Ln1 and Lg2-Ln2) expressed the chemokine receptor *Ccr6* while most cells from the minor cluster (Lg3-Ln3) did not (Figures 2A, 2B and 2E). Out of the top differentially expressed genes, we found that: Cluster 1 (Lg1-Ln1) cells expressed the IgE Fc receptor *Fcer2a* (CD23), the calcium-binding protein *Cnn3*, and the cysteine-rich protein *Lmo2*; Cluster 2 (Lg2-Ln2) cells expressed the chemokine receptor *Cxcr3*, the placenta-specific protein *Plac8*, the cystatin *Cst3* and the integrin *Itga4;* Cluster 3 (Lg3-Ln3) cells expressed the membrane-spanning 4-domains *Ms4a6c*, the transcription factor *Nfatc1*, and the zinc finger and BTB domain-containing protein *Zbtb32* (Figures 2A, 2B and 2E). Importantly, we were able to distinguish previously unreported populations of MBCs in lungs and lymph nodes by flow cytometry using the cluster-identifying markers Ccr6 and Cxcr3: Ccr6^+^Cxcr3^+^, Ccr6^+^Cxcr3^-^ and Ccr6^-^Cxcr3^-^ (Figures 2G-2H).

In spleen, MBCs from the major cluster Sp1 expressed *Ccr6, Lmo2, Fcer2a* and *Cxcr3*, a gene expression pattern associated with follicular MBCs (Figures 2C and 2E). Cells from cluster Sp2 were characterized by high levels of the Krüppel-like factor *Klf2*, the complement decay-accelerating factor *Cd55*, and the cytoskeletal protein Vimentin *Vim*, resembling transitional MBCs (Figures 2C and 2E). Cells from cluster Sp3 expressed the tyrosine kinase *Lck, Nfatc1, Ms4a6c*, and *Zbtb32*, a similar gene pattern to the one observed in B1 cells (Figures 2C and 2E). Cells from cluster Sp4 expressed high levels of the complement receptor *Cr2* (CD21) and the sphingosine 1-phosphate receptors *S1pr1* and *S1pr3*, resembling marginal zone MBCs (Figures 2C and 2E). Lastly, cells from the smallest cluster Sp5 expressed the transcription factor *Tbx21* (T-bet) and the integrin *Itgax* (Cd11c), resembling age-associated MBCs (Figures 2C and 2E) (Riedel et al., 2020). Notably, we could segregate follicular, transitional, B1-like, marginal zone and age-associated MBCs by flow cytometry based on the expression of the representative surface markers CD11c, Ccr6, CD21, CD55 and CD24 (Figure 2I). This data is in line with two potential scenarios: i) splenic B cells with diverse origins and developmental stages participate in the memory response to influenza infection; ii) MBCs acquire distinct phenotypes upon exiting germinal centers due to environmental factors. Subsequent B cell receptor (BCR) analysis presented in Figure 4 is consistent with the first model.

**Figure 3.**
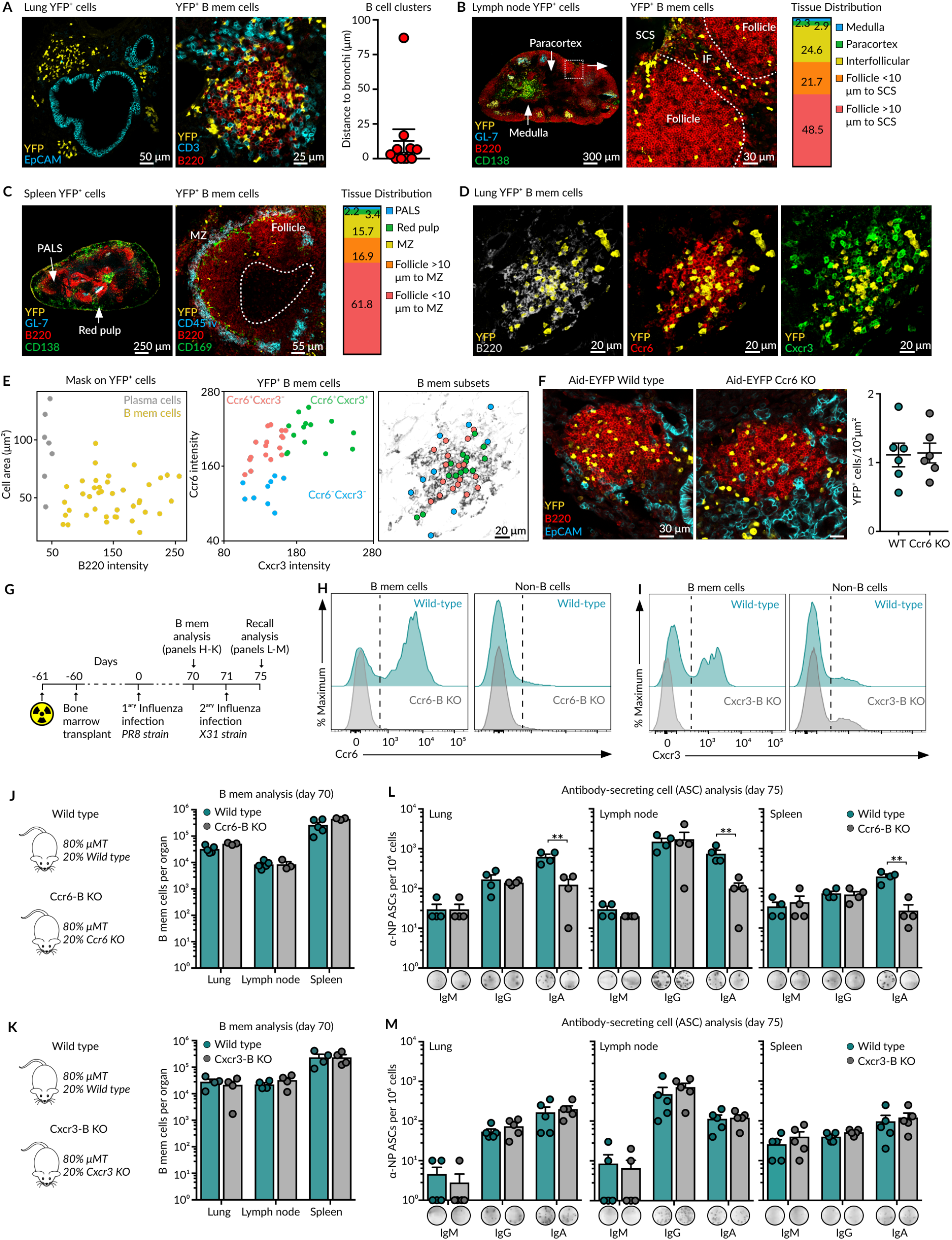
Spatial positioning of MBCs. **(A)** Confocal microscopy images of lung sections from Aid-EYFP mice treated as in Figure 1A. Sections were stained with antibodies against YFP (yellow), Epcam (cyan), B220 (red) and CD3 (cyan). Quantification displays the minimal distance of B cell clusters to the EpCAM^+^ epithelial cells. Dots represent individual B cell clusters. **(B)** Confocal microscopy images of mediastinal lymph node sections from Aid-EYFP mice treated as in Figure 1A. Sections were stained with antibodies against YFP (yellow), B220 (red), GL7 (cyan) and CD138 (green). Bar chart shows the proportion of MBCs residing in the indicated lymph node zones. **(C)** Confocal microscopy images of spleen sections from Aid-EYFP mice treated as in Figure 1A. Sections were stained with antibodies against YFP (yellow), B220 (red) and either CD138 (green) plus GL7 (cyan) or CD169 (green). CD45 (cyan) was injected i.v. in some mice to visualize the marginal zone. Bar chart shows the proportion of MBCs residing in the indicated spleen zones. **(D)** Confocal microscopy images of lung sections stained with antibodies against YFP (yellow), B220 (grey), Ccr6 (red) and Cxcr3 (green). **(E)** Dot plot showing the cell area and B220 fluorescence intensity for detected YFP^+^ surfaces in Figure 1D (left). Dot plot showing Ccr6 and Cxcr3 fluorescence intensity for individual YFP^+^ MBCs (center). X and Y positions of Ccr6^-^Cxcr3^-^ (blue), Ccr6^+^Cxcr3^-^ (red) and Ccr6^+^Cxcr3^+^ (green) MBC subsets laid out in the B220 representation (right). **(F)** Confocal microscopy images of lung sections from wild type and Ccr6-deficient Aid-EYFP mice treated as in Figure 1A. Sections were stained with antibodies against YFP (yellow), Epcam (cyan) and B220 (red). Quantification displays the density of YFP^+^ B cells in B cell clusters. Dots represent individual B cell clusters. **(G)** Overview of the strategy to generate mixed bone marrow chimeras and subsequent influenza infections. **(H-I)** Representative histograms showing Ccr6 (H) and Cxcr3 (I) expression in lung memory B and non-B cells from wild type and chemokine-deficient mixed bone marrow chimeras. **(J-K)** Quantification of MBC numbers in lungs, lymph nodes and spleen after 70 days of primary influenza infection in Ccr6-deficient (J) and Cxcr3-deficient (K) mixed bone marrow chimeras. **(L-M)** Enumeration of IgM, IgG and IgA antibody-secreting cells (ASCs) measured by ELISPOT in lungs, lymph nodes and spleen after 4 days of secondary influenza infection in Ccr6-deficient (L) and Cxcr3-deficient (M) mixed bone marrow chimeras. Quantification of one representative experiment is shown in the bar charts; each dot represents one mouse. In all panels: mean ± s.e.m, t test: **p<0.01.

**Figure 4.**
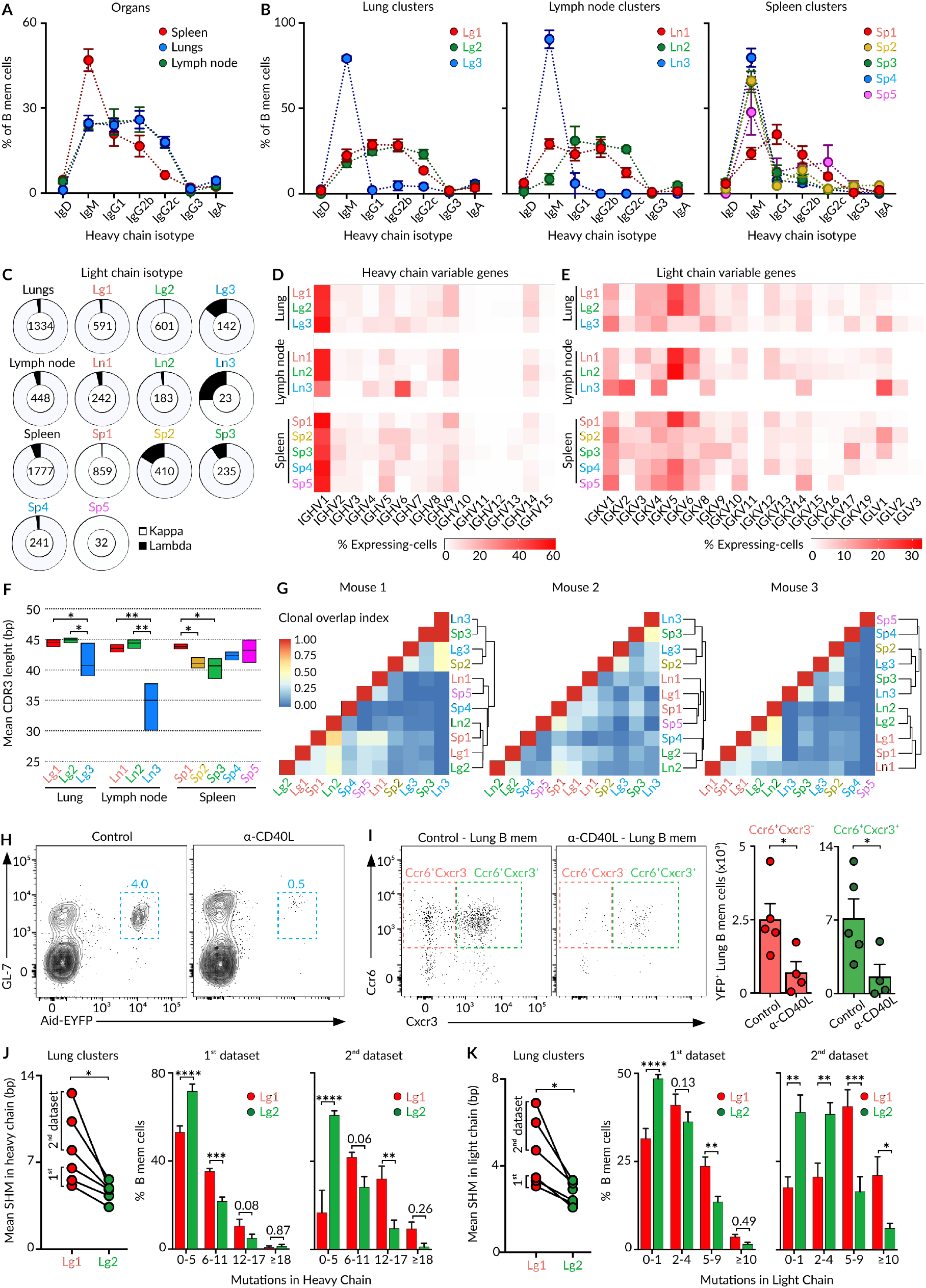
MBC progenitors. **(A-B)** Quantification of IgH isotype-class usage by MBCs according to organ (A) and tissue cluster (B) from the 1^st^ scRNA-seq dataset. **(C)** Quantification of IgK and IgL usage by MBC clusters across organs. **(D-E)** Heatmap of IgH (D) and IgK (E) variable gene frequency in individual MBC clusters. **(F)** Mean CDR3 length in IgH for indicated MBC clusters, two-way Anova test. **(G)** Szymkiewicz-Simpson similarity matrices in individual mice for pairwise comparisons of clonal overlaps among clusters and tissues. Similarity trees show hierarchical clustering analysis. **(H)** Flow cytometry plots displaying YFP^+^ germinal center B cells (previously gated on CD19^+^ cells) in mediastinal lymph nodes of Aid-EYFP animals treated as in Figure 1A. In addition, one group of mice was treated with anti-CD40L on day 6, 8 and 10. **(I)** Flow cytometry plots showing lung Ccr6^+^Cxcr3^-^ and Ccr6^+^Cxcr3^+^ MBC subsets in the CD19^+^YFP^+^CD38^+^GL7^-^ population in mice treated as in Figure 1A plus/minus administration of blocking anti-CD40L. Bar charts show the quantification of one representative experiment. Each dot represents one mouse, mean ± s.e.m, t-test. **(J-K)** Quantification of mean somatic hypermutation (SMH) levels in the variable gene of heavy (J) and light (K) chain for MBCs in clusters Lg1 and Lg2 in 1^st^ and 2^nd^ datasets, paired t-test (left). Bar charts showing the distribution of Lg1 and Lg2 cluster cells according to the levels of SHM, mean ± s.e.m, one-way Anova test (right). *p<0.05, **p<0.01, ***p<0.001 and ****p<0.0001.

It is relevant to report that in our initial analysis, we observed a fourth cluster of cells in the lungs with high expression of immediate-early response genes and heat-shock protein genes (*Hspa1a, Hspa1b, Dnajb1, Jun*) that was later removed for further analysis (Extended Figure 2A). This gene signature has been previously reported in other cell types after tissue dissociation with collagenase at 37°C (Adam et al., 2017; O’Flanagan et al., 2019). To assess if the presence of this fourth cluster was due to the dissociation protocol, we repeated the scRNA-seq experiment; this time around we dissociated lungs mechanically at 4°C rather than enzymatically. Although the number of cells recovered was significantly lower, we observed that this “stressed” subpopulation disappeared, confirming the artifactual origin of this phenomenon (Extended Figure 2B). Importantly, in this second data-set we were able to identify clusters of MBCs with gene expression patterns similar to the ones obtained in our first data-set, providing further evidence for intra and inter-organ heterogeneity of MBCs during viral infection (Extended Figures 2B-2D).

### MBC subsets colonize the lung peribronchial niche upon influenza infection

Our single-cell RNA-seq and flow cytometry data revealed the co-existence of discrete subsets of MBCs in the lung mucosa. As these subsets expressed contrasting patterns of Ccr6 and Cxcr3 receptors, we wondered if they exhibit differential micro-anatomic distributions. To gain insight into their spatial positioning, we infected Aid-EYFP mice with influenza virus and harvested lungs after 70 days. We stained tissue sections with specific antibodies and analyzed them by confocal microscopy. We found that YFP^+^ MBCs assembled in discrete B220^+^ clusters composed of hundreds of cells. These clusters were found in areas surrounding EpCAM^+^ epithelial cells, indicating that MBCs preferentially locate in peribronchial areas (Figure 3A). We further extended the analysis to lymphoid organs by tracking YFP^+^ cells devoid of germinal center (GL-7) and plasma cell (CD138) markers. In lymph nodes, MBCs preferentially accumulated in follicles and interfollicular areas but rarely in the paracortex or medulla. Strikingly, a third of follicular MBCs were found within a 10μm distance from the subcapsular sinus (Figure 3B). In spleen, YFP^+^ MBCs were mostly found in follicles, in close association with the marginal zone (Figure 3C). Altogether, these results show that MBCs remain positioned at sites of antigen entry after resolution of infection, a strategy that facilitates a fast encounter of pathogens upon re-challenge.

To visualize individual MBC subsets in the lungs, we stained tissue sections with antibodies against YFP, B220, Ccr6 and Cxcr3. We observed a wide range of Ccr6 and Cxcr3 expression among YFP^+^ cells (Figure 3D). We then generated a mask on YFP^+^ cells, excluded plasma cells based on their size and B220 expression, and measured Ccr6 and Cxcr3 fluorescence on individual YFP^+^ cells. By plotting Cxcr3 versus Ccr6 fluorescence intensity, we observed a cell distribution pattern that closely resembled the one obtained by flow cytometry (Figures 2G and 3E). Interestingly, when we tracked each subset back into space, we found that cells from the two main Ccr6^+^ subsets localized in the center of B cell clusters while cells lacking Ccr6 were excluded to the peripheral areas (Figure 3E). Due to this marked differential positioning, we investigated if Ccr6 was required by MBCs to access peribronchial B cell clusters. To this end, we generated Aid-EYFP mice lacking Ccr6 expression, infected them with influenza virus and analyzed lung sections at day 70 of infection. We found similar numbers of YFP^+^ MBCs within B cell clusters in wild type and Ccr6-deficient mice, indicating that Ccr6, *per se*, is dispensable to enter the peribronchial niche (Figure 3F). Altogether, our analysis revealed a new layer of organization at the core of peribronchial B cell clusters, with MBCs segregating into discrete regions.

To further test if Ccr6 and Cxcr3 chemokine receptors play a role in the retention of MBCs in the lung mucosa or lymphoid organs, we generated chimeric mice by lethally irradiating µMT animals and injecting a mixture of 80% µMT bone marrow and 20% of either wild type, Ccr6- or Cxcr3-deficient bone marrow (Figure 3G). As µMT mice lack mature B cells, chemokine-knockout chimeras harbored Ccr6 or Cxcr3-deficient B cells in an environment consisting of mostly wild type cells (Figures 3H-3I). After 8 weeks of reconstitution, we infected chimeric mice with 5 PFU of H1N1 PR8 influenza virus and sacrificed them 70 days after for analysis. Surprisingly, we observed comparable numbers of CD19^+^IgD^-^CD38^+^GL-7^-^ MBCs among wild type, Ccr6 and Cxcr3-deficient chimeras, indicating that these receptors are dispensable for MBC maintenance (Figures 3J-3K). We then assessed if Ccr6 and Cxcr3 are required for the memory response during secondary influenza infection. To test this, we infected chimeric mice with influenza virus H1N1 PR8 and after 70 days we challenged them with 5.10^4^ PFU of influenza H3N2 X31 strain. Four days after re-infection, we measured the formation of IgM, IgG and IgA NP-specific antibody-secreting cells (ASCs) by ELISpot. We observed a 10-fold decrease in the formation of IgA ASCs in Ccr6-deficient chimeras compared to wild type counterparts, while the numbers of IgM and IgG ASCs were similar among both groups (Figure 3L). This defect was observed in lungs, lymph nodes and spleen, suggesting a general mechanism for Ccr6 in IgA responses across organs. In contrast, we observed comparable numbers of ASCs in wild type and Cxcr3-deficient chimeras, indicating that Cxcr3 is dispensable for memory responses during influenza infection (Figure 3M). Overall, our results show that Ccr6, but not Cxcr3, is required for recall of humoral responses to influenza virus.

### Lung MBC subsets undergo class-switching and somatic hypermutation in germinal centers

Having shown that discrete subsets of MBCs with divergent transcriptional programs and tissue distribution co-exist upon resolution of influenza infection, we sought to investigate if the subsets arise from common or distinct progenitors. To achieve this, we produced and sequenced single-cell BCR-seq libraries from the same cells analyzed by droplet-based scRNA-seq. Analysis of the constant heavy chain (IgH) revealed that MBCs residing in lungs and lymph nodes displayed extensive class-switching towards IgG isotypes, such as IgG1, IgG2b and IgG2c, while the majority of splenic memory cells expressed IgM (Figure 4A). To assess if this difference was due to the presence of different B cell subpopulations across organs, we analyzed isotype expression individually in each cell cluster. In lungs and lymph nodes, the two major Ccr6^+^ clusters (Lg1-Ln1 and Lg2-Ln2) showed considerable class-switching towards IgG isotypes while the minor Ccr6^-^ cluster (Lg3-Ln3) was almost exclusively composed of IgM^+^ cells (Figure 4B). In spleen, the major follicular MBC cluster (Sp1) exhibited substantial class switching towards IgG isotypes while transitional (Sp2), B1-like (Sp3), marginal zone (Sp4) and ABC (Sp5) MBCs were mainly IgM^+^ (Figure 4B). Concerning the constant light chain, we observed that Kappa (IgK) was the most commonly expressed among MBCs regardless of the cell cluster and organ of origin (Figure 4C). Yet, we found unusual high numbers of Lamba^+^ cells in the minor IgM^+^Ccr6^-^ cluster (Lg3-Ln3) from lungs/lymph nodes and in transitional (Sp2) and B1-like (Sp3) MBCs from spleen (Figure 4C). Altogether, these results unveiled that the MBC pool displays pronounced intra- and inter-organ differences on isotype-class usage.

The marked heterogeneity on isotype-classes led us to investigate the magnitude of BCR diversification driven by the differential use of variable genes (IgHV, IgKV and IgLV) and the length of the complementary-determining region 3 (CDR3) within each subset. We observed that the two major class-switched Ccr6^+^ clusters (Lg1-Ln1 and Lg2-Ln2) in lungs/lymph nodes and the follicular (Sp1), marginal zone (Sp4) and ABC (Sp5) clusters in spleen, displayed high usage of the IgHV1, 5, 9, 14 and IgKV5, 6 families (Figures 4D and 4E). By contrast, the minor IgM^+^Ccr6^-^ clusters (Lg3-Ln3) from lungs/lymph nodes and the transitional (Sp2) and B1-like (Sp3) clusters from spleen showed a completely different pattern of expression: high usage of IgHV1, 4, 6; IgKV1, 2, 4, 8, 14 and IgLV1 (Figures 4D and 4E). Remarkably, these clusters further exhibited a significantly lower length of the IgH CDR3 when compared to clusters from the same organ (Figure 4F). These results demonstrate that MBC subsets are skewed towards divergent families of variable genes, in line with the notion that certain memory subsets may originate from different naïve precursors. To further test this idea, we defined clonotypes and measured clonal overlaps in MBC clusters. We detected significant overlap of clonal BCR repertoires among lung/lymph node clusters containing class-switched Ccr6^+^ cells and the splenic follicular cluster, Lg1-Ln1-Sp1 on one side and Lg2-Ln2-Sp1 on the other side (Figure 4G). We further detected high levels of clonal overlap among the lung/lymph node clusters enriched in IgM^+^Ccr6^-^ cells and splenic transitional and B1-like clusters (Lg3-Ln3-Sp2-Sp3) (Figure 4G). These results imply that certain memory populations are interconnected in their generation and/or maintenance across organs. Remarkably, hierarchical clustering showed that the minor “innate-like” group of IgM^+^Ccr6^-^ cells represented a separate lineage from class-switched Ccr6^+^ MBCs, providing further evidence that these populations emerge from different progenitors (Figure 4G).

The two major class-switched Ccr6^+^ memory populations identified in lungs and draining lymph nodes, display divergent transcriptional programs and can be segregated based on their Cxcr3 expression (Figure 2). To evaluate whether these two subsets were both product of germinal center reactions, we infected two groups of Aid-EYFP mice with influenza virus H1N1 and between days 6-10 we administered tamoxifen with either blocking anti-CD40L antibody or isotype control. By using this window of time, we allow early B-T cell contacts while targeting subsequent B-T follicular Helper cell interactions in germinal centers. We observed a dramatic reduction of YFP^+^ germinal center cells in mice treated with anti-CD40L antibody compared to control animals (Figure 4H). Importantly, we found that the Cxcr3^-^ and the Cxcr3^+^ subsets were markedly reduced in mice receiving anti-CD40L treatment, strongly suggesting that both MBC subsets are products of germinal centers (Figure 4I). We then measured if these two populations arose from B cell precursors that played an active role in germinal center reactions. To assess this, we measured the average rate of somatic hypermutation (SHM) in IgHV and IgKV genes in our two scRNA-seq datasets. We found that the Lg1 cluster (Cxcr3 low) displayed significantly higher average levels of SHM in the IgHV and IgKV genes than the Lg2 cluster (Cxcr3 high) (Figures 4J and 4K). When analyzing SHM in individual cells, we found that the Lg1 cluster (Cxcr3 low) is enriched in cells with high levels of mutations while the Lg2 cluster (Cxcr3 high) is enriched in cells with lower SHM levels (Figures 4J and 4K). These results indicate that the two major Ccr6^+^ class-switched MBC subsets derived from cells that underwent SHM in germinal centers, but either at dissimilar rates or for different periods of time.

### Lung MBC subsets display contrasting specificities and protective functions

We wondered whether these two transcriptionally distinct subsets of class-switched Ccr6^+^ MBCs were intrinsically biased in their differentiation fate towards plasma versus germinal center cells upon re-activation. To evaluate this, we sorted Ccr6^+^Cxcr3^-^ and Ccr6^+^Cxcr3^+^ MBC subsets from lungs and lymph node at day 70 after influenza infection and cultured them ex vivo in the presence of 40LB feeder cells, sources of BAFF and CD40-L, plus IL-21 for 3 days (Figure 5A) (Nojima et al., 2011). Under these conditions, MBCs have been shown to differentiate into plasma or germinal center cells according to cell intrinsic features (Koike et al., 2019). Flow cytometry analysis revealed that both Ccr6^+^ MBC subsets mostly differentiated into CD93^+^ plasma cells and rarely into GL-7^+^ germinal center B cells, regardless of their Cxcr3 expression or organ of origin (Figures 5B-5C). Furthermore, we detected similar IgG levels across samples, indicating that these cells produce antibodies to the same extent (Figure 5D). Unexpectedly, we found that plasma cells derived from Cxcr3^+^ MBCs produced high levels of antibodies specific for influenza NP and HA while plasma cells derived from Cxcr3^-^ MBCs produced low to undetectable levels of NP and HA-specific antibodies (Figure 5E). These results demonstrate that Cxcr3^-^ and Cxcr3^+^ MBC subsets are both intrinsically predisposed towards the plasma cell program but show profound differences in their ability to produce virus-specific antibodies upon activation. We then examined the specificity of MBC subsets for influenza antigens. To this end, we incubated lung and lymph node cell suspensions from influenza infected mice (day 70) with fluorescently labelled NP and HA. Flow cytometry analysis revealed that Cxcr3^+^ MBCs bound extensively to influenza antigens while Cxcr3^-^ cells showed minimal binding (Figures 5F-5G). Thus, the Cxcr3^+^ MBC subset is indeed enriched in antigen specific cells while the Cxcr3^-^ subset comprises cells with no apparent specificity for influenza proteins.

**Figure 5.**
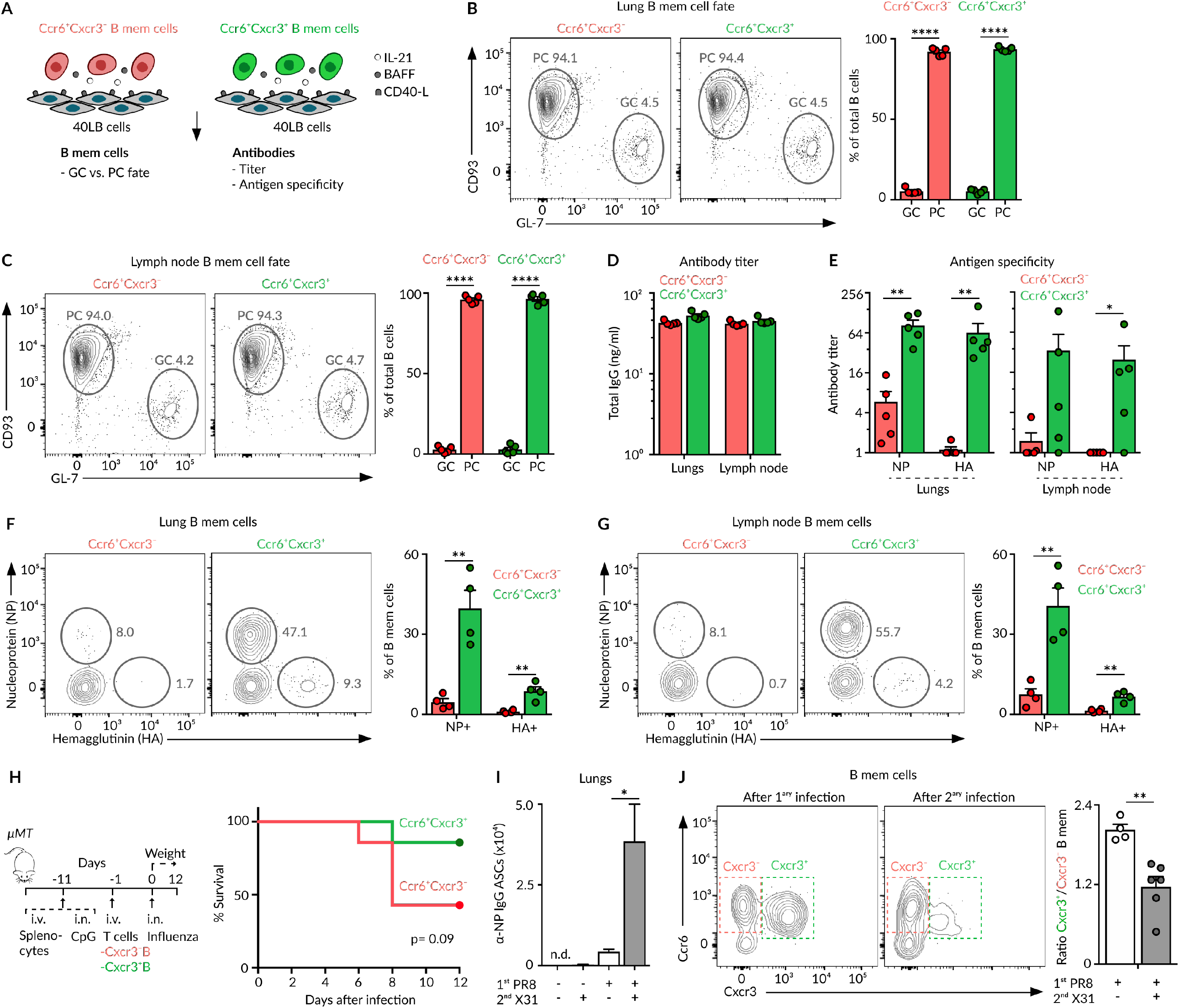
MBC fate and specificity. **(A)** Schematic overview of the 40LB-MBC co-culture in the presence of IL-21 (10ng/ml). Listed are the measured outputs. **(B-C)** Flow cytometry plots showing CD93^+^ plasma and GL7^+^ germinal center-like cells derived from either Ccr6^+^Cxcr3^-^ or Ccr6^+^Cxcr3^+^ MBCs isolated from lungs (B) and lymph nodes (C) at day 70 of influenza infection and co-cultured for 3 days with 40LB cells. **(D-E)** Total IgG levels (D), NP and HA-specific IgG titers (E) measured at day 3 in supernatants of Ccr6^+^Cxcr3^-^ and Ccr6^+^Cxcr3^+^ MBC co-cultures with 40LB cells. **(F-G)** Flow cytometry plots showing the percentage of Ccr6^+^Cxcr3^-^ or Ccr6^+^Cxcr3^+^ MBCs from lungs (F) and lymph nodes (G) of mice treated as in Figure 1A, that bind to fluorescently labelled Nucleoprotein (NP) and Hemagglutinin (HA). **(H)** Overview of the strategy to evaluate the protective capacity of Ccr6^+^Cxcr3^-^ or Ccr6^+^Cxcr3^+^ MBCs in vivo upon infection with Influenza PR8. Survival curve of mice treated as in the left scheme. **(I)** Enumeration of NP-specific IgG antibody-secreting cells measured by ELISPOT in lungs of mice treated with PBS or influenza PR8 and challenged at day 70 with PBS or Influenza X31. Analysis was performed after 4 days of secondary challenge. **(J)** Flow cytometry plots displaying Ccr6^+^Cxcr3^-^ and Ccr6^+^Cxcr3^+^ MBCs in lungs of mice infected with influenza PR8 alone (left panel) or challenged with influenza X31 after 70 days (right panel). Quantification of one representative experiment is shown in bar chart; mean ± s.e.m, each dot represents one mouse. t test: *p<0.05, **p<0.01 and ****p<0.0001.

These unforeseen results led us to investigate the involvement of these two MBC subsets in homotypic and heterosubtypic immunity to influenza infection. For the former, we took advantage of a previously described in vivo set-up (Onodera et al., 2012). We i.v. transferred naive splenocytes to µMT mice and treated them intranasally with CpG to generate bronchoalveolar lymphoid structures. After ten days, we transferred Cxcr3^-^ or Cxcr3^+^ MBCs sorted from lungs of previously PR8-infected animals together with splenic CD4^+^ T cells isolated from the same donors. We intranasally infected these animals with influenza PR8, monitored weight daily and euthanized those mice exhibiting ≥20% loss of initial mass (Figure 5H). We observed that mice receiving Cxcr3^+^ MBCs were highly protected from influenza infection while more than 50% of those receiving Cxcr3^-^ MBCs needed to be euthanized between day 6 and 8 (Figure 5H). Therefore, Cxcr3^+^ and Cxcr3^-^ subsets differ in their ability to provide homotypic protection. To investigate the ability of these subsets to intrinsically respond to re-infection with a different influenza serotype, we intranasally treated Aid-EYFP mice with PBS or influenza PR8 and gave them tamoxifen from days 6-10. At day 70, we intranasally challenged these mice with either PBS or influenza X31 and sacrificed them after 4 days. Secondary infection with X31 led to robust formation of flu-specific antibody-secreting cells (Figure 5I). These cells were mainly derived from lung MBCs, as this response was not observed in mice receiving PR8 or X31 infection alone (Figure 5I). Taking this into account, if one subset of MBCs is preferentially engaged into secondary responses, we should observe a change in subsets ratio before and after re-challenge (Allie et al., 2019). We found that the Cxcr3^+^/Cxcr3^-^ memory ratio is close to 2 after primary infection (PR8 alone) (Figure 5J). Interestingly, we observed a significant and marked decrease in the Cxcr3^+^/Cxcr3^-^ ratio upon secondary infection (PR8 + X31), indicating that Cxcr3^+^ MBCs were actively differentiating into effector cells when compared to the Cxcr3^-^ subset (Figure 5J). Altogether, these results show that influenza infection gives rise to two distinct populations of Ccr6^+^ MBCs in lungs: i) a Cxcr3^+^ subset enriched in MBCs with specificity for influenza antigens that robustly respond upon secondary challenge by secreting protective antibodies, and ii) a Cxcr3^-^ subset enriched in MBCs with no evident specificity for influenza antigens and restricted effector capacity in recall responses.

### Cxcr3 expression segregates lung MBCs into bona fide and bystander subsets

We were intrigued by the results showing that the Ccr6^+^Cxcr3^-^ MBC subset is composed of cells that underwent class-switching and somatic hypermutation but do not display evident specificity for influenza antigens. This led us to consider three hypotheses regarding their origin: i) these cells are low-affinity MBCs with undetectable antigen capture in our flow cytometry antigen-binding assay, ii) these cells were initially antigen-specific but subsequently acquired disadvantageous mutations and escaped germinal center reactions as non-specific MBCs, iii) these are non antigen-specific cells that were selected in germinal centers through alternative permissive mechanisms. While in the first two scenarios Cxcr3^-^ MBCs can share progenitors with antigen-binding cells, no clonal overlap is expected in the third possibility. To discern among these contrasting situations, we performed single-cell index sorting of lung YFP^+^ MBCs from 3 mice at day 70 of influenza infection. For each cell, we recorded the information of NP/HA antigen-binding and the expression levels of surface Ccr6/Cxcr3 receptors. We then performed FACS-Based 5-Prime End (FB5P) single-cell RNA-seq on sorted cells for the integrative analysis of transcriptome and BCR repertoire (Attaf et al., 2020). After processing and filtering, we retained 983 good quality cells with information on heavy and light chains for subsequent analysis. In this dataset, most cells that bound NP/HA antigens were part of the Ccr6^+^Cxcr3^+^ subset, reinforcing our previous results (Figure 6A). We further measured the expression of cluster-identifying genes from our first scRNA-seq datasets in the different MBC subsets (Figure 2E and 6B). We found that the Ccr6^+^Cxcr3^-^ subset express high levels of Lg1 cluster genes (*Fcer2a, Cnn3, Lmo2*), the Ccr6^+^Cxcr3^+^ subset express high levels of Lg2 cluster genes (*Cst3, Itga4*) and the minor Ccr6^-^Cxcr3^-^ subset is characterized by the expression of Lg3 cluster genes (*Ms4a6c, Zbtb32*) (Figure 6B). These results demonstrate that the segregation of MBCs based on the expression of Ccr6 and Cxcr3 is an accurate representation of the transcriptionally divergent populations identified by scRNA-seq.

**Figure 6.**
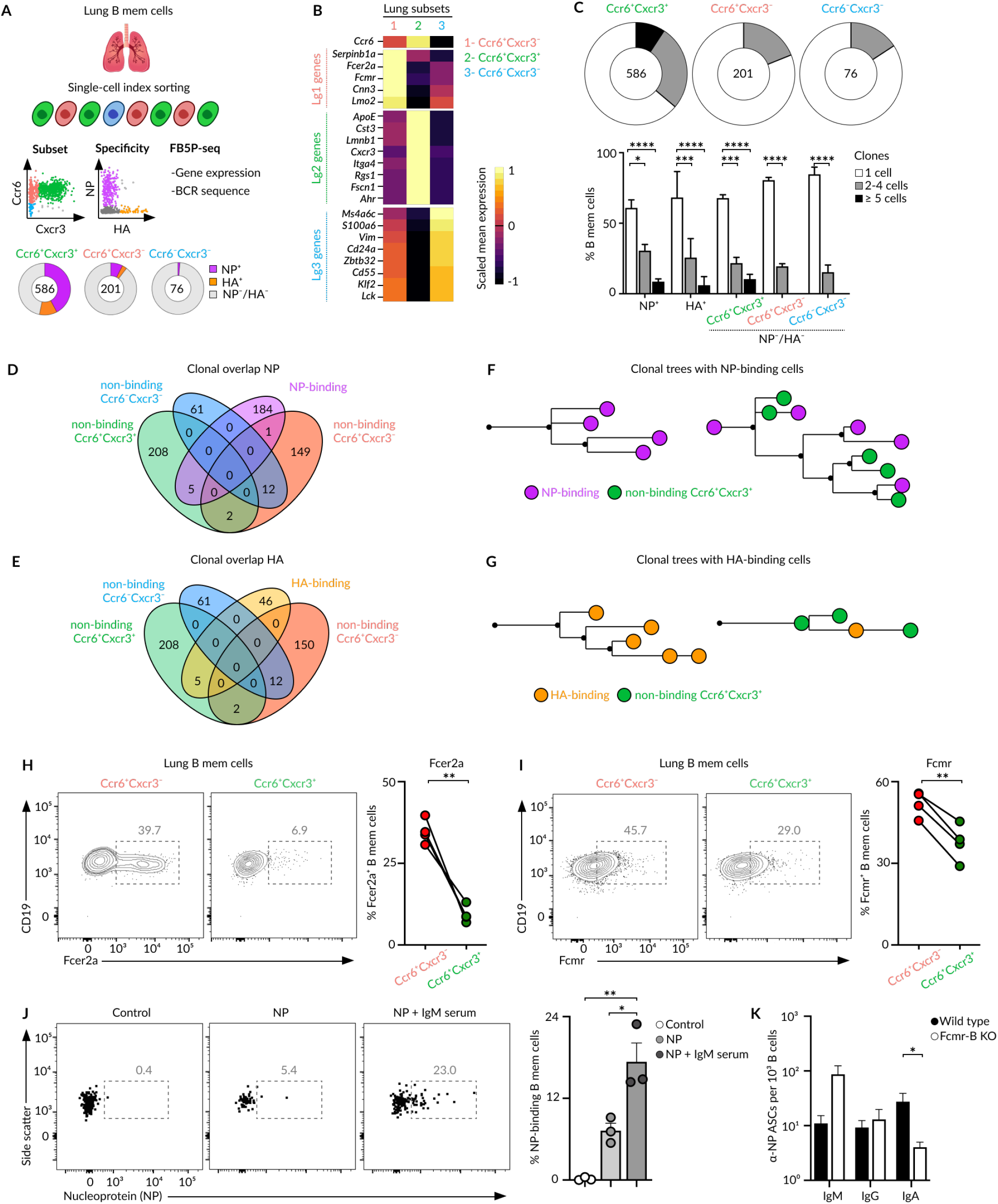
Bona fide and bystander MBCs. **(A)** Overview of the index cell sorting and FB5P-seq experimental workflow. Information of Ccr6/Cxcr3 expression and HA/NP binding was recorded for each lung MBC sorted. Quantification of HA/NP binding cells in different memory subsets is shown in pie charts. **(B)** Heatmap showing the expression of marker genes from Lg1, Lg2 and Lg3 clusters (10X dataset) by Ccr6^+^Cxcr3^-^, Ccr6^+^Cxcr3^+^ and Ccr6^-^Cxcr3^-^ subsets. Colour: Scaled mean expression. **(C)** Pie charts displaying the percentage of cells comprising single-cell, 2-4 cell, or ≥4 cell clones for each subset. Bar charts showing clonal size in NP/HA binding and non-binding memory cells, two-way Anova test. **(D-E)** Venn diagrams showing the clonal overlap among non-binding cells from Ccr6^+^Cxcr3^-^, Ccr6^+^Cxcr3^+^ and Ccr6^-^Cxcr3^-^ subsets and NP-binding (D) or HA-binding (E) cells. **(F-G)** Trees showing phylogenetic relationships of IgH and IgK sequences from clones containing NP-binding (F) and HA-binding (G) cells. **(H-I)** Flow cytometry plots showing the expression of *Fcer2a* (H) and *Fcmr* (I) by Ccr6^+^Cxcr3^-^ and Ccr6^+^Cxcr3^+^ MBC subsets, paired t-test. **(J)** Flow cytometry plots showing NP binding to Ccr6^+^Cxcr3^-^ lung MBCs after incubation with media, NP or NP-IgM from immune sera, t-test. **(K)** Enumeration of IgM, IgG and IgA ASCs measured by ELISPOT in lungs of chimeric mice with a wild type or Fcmr-deficient B cell compartment. Mice were infected with Influenza PR8, challenged with influenza X31 after 40 days and sacrificed 4 days later, Anova test. In all panels, each dot represents one mouse, mean ± s.e.m, *p<0.01.*p<0.05, ***p<0.001 and ****p<0.0001.

We then examined to which extent the Ccr6^+^Cxcr3^+^, Ccr6^+^Cxcr3^-^ and Ccr6^-^Cxcr3^-^ subsets were clonally expanded. Regardless of the subset, we found that more than 60% of MBCs were constituted by single-cell clones, in line with recent observations from the Nussenzweig lab (Figure 6C) (Viant et al., 2020). Furthermore, of those cells that were clonally expanded, the vast majority formed small clones of 2-4 cells. Only 6% of MBCs formed largely expanded clones (>4 cells), a phenomenon that seemed to be independent of antigen-binding as similar clonal size distributions were observed in HA/NP-binding and non-binding cells (Figure 6C). Despite the limited number of expanded clones, we moved on to evaluate the overlaps among NP and HA-binding MBCs with non-binding cells from the different subsets. Remarkably, we found notable overlaps among NP/HA-binding cells with non-binding cells from the Ccr6^+^Cxcr3^+^ subset; non significant overlaps were detected with Ccr6^+^Cxcr3^-^ and Ccr6^-^Cxcr3^-^ subsets (Figures 6D-6E). Furthermore, phylogenetic trees based on heavy and light chain nucleotide sequences show that clones comprising NP/HA-specific cells were either entirely composed of antigen-binding cells or contained several non-binding cells; regardless of the situation, these cells exclusively corresponded to the Ccr6^+^Cxcr3^+^ subset (Figures 6F-6G). Overall, our results are in line with scenario number (iii).

To further strengthen this notion, we aimed to explore cells expressing influenza-associated BCRs in a larger dataset. To achieve this, we firstly identified common BCR genes used by NP and HA-binding cells across all mice analyzed. We found that a large fraction of NP-binding cells uses IGHVs 9-3 or 9-4 and IGKVs 5-48, 5-37 or 5-43 genes, indicating that the NP-specific BCR repertoire is in part composed of “public” IgHV / IgKV combinations (Extended Figure 6A). Although we detected common usage of IGHVs 14-2/1-63 and IGKV 4-72 by HA-specific B cells, numbers were not enough to define HA-associated BCRs with high confidence (Extended Figure 6B). Then, we used VH 9-3/4 and VK 5-48/37/43 variable gene usage to identify putative NP-specific B cells across lungs, lymph nodes and spleen in our large 10x dataset. We detected 376 VH 9-3/4^+^ VK 5-48/37/43^+^ cells among the 3559 cells with paired IgH+IgKL sequences. Remarkably, in all organs and mice analyzed, we observed higher numbers of *Cxcr3*-expressing cells in the putative NP-specific population than in the one expressing alternative BCR gene combinations (Extended Figure 6C).

Altogether, the simultaneous analysis of antigen-binding, surface marker expression and B cell repertoire at the single cell level revealed that Ccr6^+^Cxcr3^+^ and Ccr6^+^Cxcr3^-^ populations represent discrete subsets of MBCs with divergent origins. The Cxcr3^+^ subset is composed of antigen-specific *bona fide* MBCs that share common progenitors with low-affinity cells. In contrast, the Cxcr3^-^ subset is enriched in *bystander* MBCs, with no apparent specificity for influenza antigens and originating from highly permissive selection mechanisms in germinal centers.

Remarkably, FB5P-seq transcriptomic analysis showed that *bystander* Cxcr3^-^ MBCs express higher levels of Fc receptors for IgE and IgM (*Fcer2a* and *Fcmr*) than Cxcr3^+^ *bona fide* MBCs (Figure 6B). We confirmed this observation at the protein level by flow cytometry (Figures 6H-6I). This led us to investigate if *bystander* MBCs could bind immune complexes despite their lack of specificity for influenza antigens, a phenomenon that would increase antigen density in the proximity of *bonafide* MBCs. We found that incubation of lung MBCs with influenza-IgM immune complexes led to significant binding to *bystander* MBCs (Figure 6J). To assess the contribution of this axis on the magnitude of memory responses, we generated mouse chimeras lacking *Fcmr* specifically on the B cell compartment. Chimeric mice were infected with influenza H1N1 and re-challenged with influenza H3N2 at day 40. We found a profound reduction in the generation of influenza-specific IgA ASCs in *Fcmr*-deficient mice, providing evidence for a role of the *Fcmr* axis on MBC responses (Figure 6K).

### SARS-CoV-2 infection generates bona fide and bystander MBC subsets

We were interested to find out whether the establishment of discrete subsets of MBCs in the respiratory environment with contrasting antigen specificities is a general feature of humoral responses to airborne pathogens. To address this question, we intranasally infected K18-hACE2 mice, which express the human angiotensin-converting enzyme 2 receptor under the promoter of cytokeratin 18, with 2.10^3^ PFU of SARS-CoV-2 (McCray et al., 2007). We stained lung sections with antibodies against EpCAM and SARS-CoV-2 NP to track the kinetics of infection by confocal microscopy. We detected foci of SARS NP in peribronchial areas as early as 2 days after infection while the infection rapidly spread towards deeper areas of the lung parenchyma by day 5 (Figure 7A). Remarkably, we did not detect SARS-NP in lung tissues by day 70 of infection, in clear contrast to what has been reported in human gut biopsies, where SARS NP is detected even after 3 months of COVID-19 symptom onset (Gaebler et al., 2021). Furthermore, no effective infection was observed in wild type C57BL/6 mice, confirming previous findings that SARS-CoV-2 does not use mouse ACE2 to infect cells (Figure 7B) (Zhou et al., 2020).

**Figure 7.**
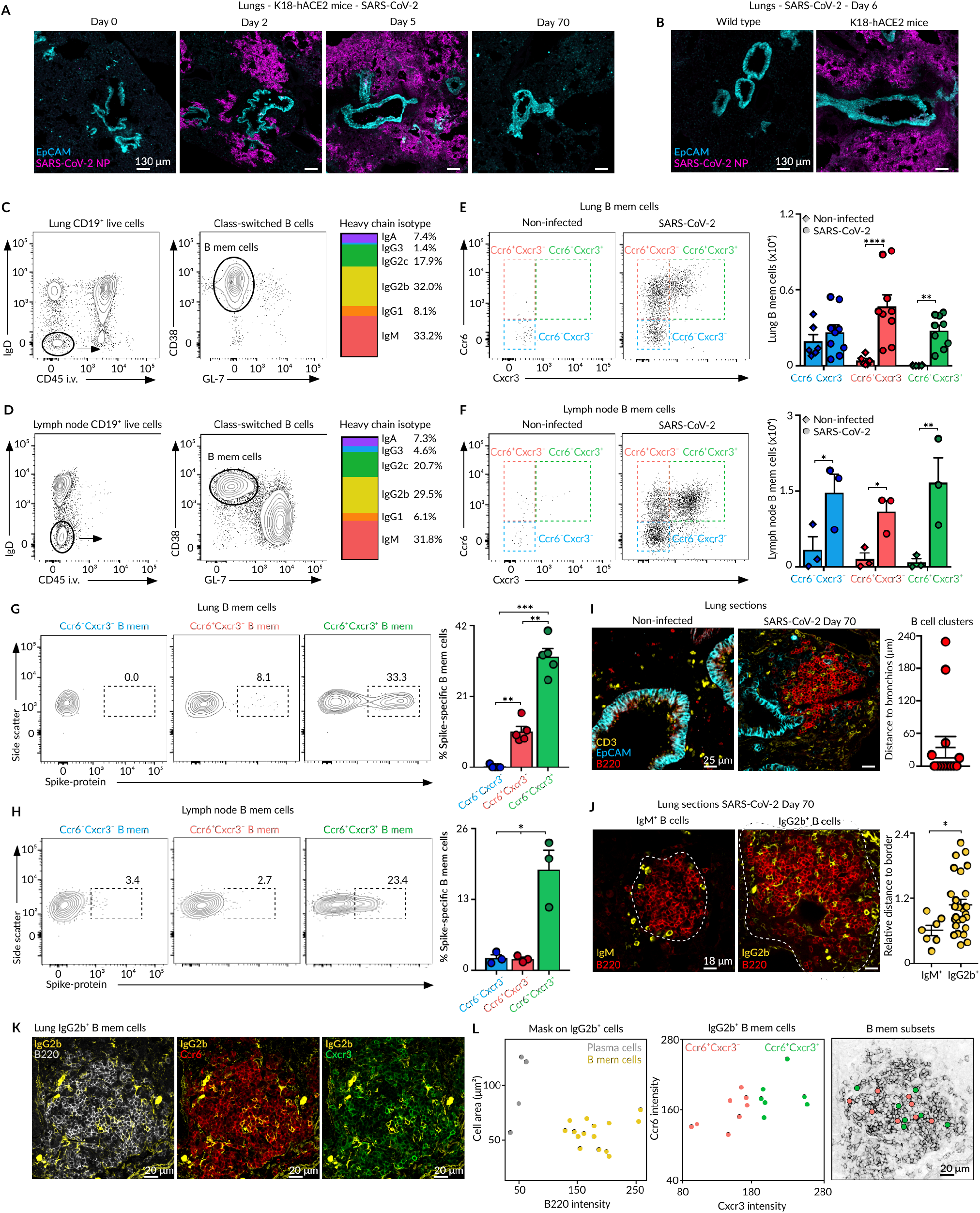
MBC subsets in SARS-CoV-2 infection. **(A-B)** Confocal microscopy images of lung sections from K18-hACE2 (A-B) or wild type (B) mice infected with 2.10^3^ PFU of SARS-CoV-2 and analyzed at indicated time points. Sections were stained with antibodies against EpCAM (cyan) and SARS-CoV-2 NP (magenta). **(C-D)** Flow cytometry plots showing lung (C) and lymph node (D) class-switched resident B cells (left, CD19^+^IgD^-^CD45i.v.^-^) and MBCs (middle, CD38^+^GL-7^-^) from K18-hACE2 infected as in (A) and analyzed at day 70. Multicolor bar (right) shows the proportion of MBCs expressing different IgH isotypes. **(E-F)** Dot plots showing the presence of Ccr6^-^ Cxcr3^-^, Ccr6^+^ Cxcr3^-^ and Ccr6^+^ Cxcr3^+^ MBC subsets in lungs (E) and lymph nodes (F) of K18-hACE2 mice left untreated or infected as in (A). **(G-H)** Flow cytometry plots showing the percentage of MBC subsets that bind to PE labelled Spike protein on lungs (G) and mediastinal lymph nodes (H). **(I-J)** Confocal microscopy images of lung sections from K18-hACE2 mice left untreated or infected as in (A) and analyzed at day 70. Sections were stained with antibodies against EpCAM (cyan), CD3 (yellow) and B220 (red) (I) or IgM (yellow), IgG2b (yellow) and B220 (red) (J). Quantification in (I) displays the minimal distance of B cell clusters to the EpCAM^+^ epithelial cells; dots represent individual B cell clusters. Quantification in (J) displays the minimal distance (nm) of individual IgM^+^ or IgG2b^+^ cells to the B cell cluster border divided by the area (µm^2^) of the cluster; dots represent individual B cells. **(K)** Confocal microscopy images of lung sections from K18-hACE2 mice treated as in (A) and stained with antibodies against IgG2b (yellow), B220 (grey), Ccr6 (red) and Cxcr3 (green). **(L)** Dot plot showing the area and B220 fluorescence intensity for detected IgG2b^+^ surfaces in Figure 7K (left). Dot plot showing Ccr6 and Cxcr3 fluorescence intensity for individual IgG2b^+^ MBCs (center). X and Y positions of Ccr6^+^Cxcr3^-^ (red) and Ccr6^+^Cxcr3^+^ (green) MBC subsets laid out in the B220 representation (right). In panels E to H, bar charts show the quantification of one representative experiment out of three, mean ± s.e.m. Each dot represents one mouse. (E-F) Two-way Anova test, (G-H) One-way Anova test. *p<0.05, **p<0.01, ***p<0.001 and ****p<0.0001.

Having set up this acute SARS-CoV-2 infection model in K18-hACE2 mice, we proceeded to analyze MBC populations by flow cytometry 70 days after infection. We administered FITC-labelled anti-CD45 antibody intravenously 5 minutes before sacrifice to exclude circulatory B cells. We observed accumulation of class-switched MBCs (CD19^+^ CD45i.v.^-^ IgD ^-^CD38^+^ GL-7^-^) in both lungs and mediastinal lymph nodes and the persistence of long-lived germinal center reactions (CD38^-^ GL-7^+^) exclusively in lymph nodes (Figures 7C and 7D). MBCs showed extensive class-switching towards IgG subclasses 2b and 2c (Figures 7C and 7D). Remarkably, we distinguished three discrete populations of MBCs across lungs and lymph nodes based on their differential expression of Ccr6 and Cxcr3 receptors (Figures 7E and 7F). The Ccr6^+^Cxcr3^+^ cells extensively bound fluorescently labelled SARS-CoV-2 spike protein, indicating that this population represents the bona fide subset enriched in virus-specific cells (Figures 7G and 7H). In contrast, the Ccr6^+^Cxcr3^-^ and Ccr6^-^Cxcr3^-^ cells did not show substantial binding to spike protein, indicating that these populations represent bystander and innate-like subsets respectively. To be noted, a significant number of lung-resident memory B and T cells was still observed in mice infected with low doses of SARS-CoV-2 virus, even in those with no apparent weight loss (Extended Figure 7). These results show that asymptomatic animals can still mount lung memory B and T cell responses to SARS-CoV-2.

We finally analyzed the spatial distribution of MBCs in lungs of SARS-CoV-2 infected animals by confocal imaging. As in this case we cannot rely on YFP expression, we stained lung sections with antibodies against B220, IgM and IgG2b to track MBCs. We found that SARS-CoV-2 infection triggers the formation of B cell clusters that accumulate in peribronchial areas (Figure 7I). Interestingly, while IgM^+^ cells were restricted to the periphery of B cell clusters, IgG2b^+^ cells were largely confined in the center (Figure 7J). Analysis of Ccr6 and Cxcr3 expression in individual IgG2b^+^ cells, revealed that both Ccr6^+^Cxcr3^+^ and Ccr6^+^Cxcr3^-^ populations were equally distributed inside the B cell clusters (Figures 7K-7L).

Altogether, our results show that SARS-CoV-2 infection, as observed with influenza virus, triggers the accumulation of bona fide, bystander and innate-like MBCs in the respiratory niche with marked differences on antigen-specificity, tissue localization and chemokine receptor expression. These results unveil common MBC mechanisms elicited during infection with diverse families of respiratory viruses.

## Discussion

Several studies have shown that the MBC pool is composed of discrete cell subsets that coexist upon immunization and that bear a differential ability to seed germinal center reactions or form antibody-secreting cells upon re-challenge (Weisel and Shlomchik, 2017). Some research lines demonstrated that MBC fate is associated with antibody isotype expression: while IgM^+^ MBCs are more predisposed to re-enter germinal center reactions, those expressing IgG are more prone to acquire an antibody-secreting phenotype upon re-challenge (Dogan et al., 2009; Kometani et al., 2013; Lutz et al., 2015; Pape et al., 2011). However, antibody isotype expression do not always draw a clear-cut line, as class-switched cells were also shown to actively remodel their BCR within secondary germinal center reactions (McHeyzer-Williams et al., 2015). Work from Shlomchik and collaborators showed that subcategorization of MBCs according to the expression of CD80 and PD-L2, independently of antibody isotype, identify cell subsets with distinct developmental kinetics and functions upon rechallenge (Anderson et al., 2007; Weisel et al., 2016; Zuccarino-Catania et al., 2014). This differential ability to commit to the germinal center or plasma cell program by CD80-high and CD80-low memory subsets is associated with the unequal amount of CD40L help received (Koike et al., 2019). As most of these studies were performed in the context of protein immunization, it remained unknown whether distinct subsets of MBCs coexist upon resolution of infection. Here, we used unsupervised transcriptomic analysis to unveil the level of MBC heterogeneity in lungs, mediastinal lymph node and spleen upon respiratory influenza virus infection.

Our study revealed that the MBC pool displays previously unrecognized high levels of heterogeneity across secondary lymphoid organs. For instance, the spleen constitutes a niche for at least five subsets of MBCs that cohabitate this organ upon resolution of influenza infection: follicular, transitional, B1-like, marginal zone and age-associated (ABC) MBCs. While the follicular population exhibits certain levels of class-switching, the other subsets are highly enriched in IgM^+^ MBCs. In line with recent studies, these results show that various splenic B cell populations can participate in the development of a memory response (Berry et al., 2020; Krishnamurty et al., 2016; Riedel et al., 2020; Stone et al., 2019). Future work is required to establish the contribution of each of these splenic subsets to secondary immune responses upon re-challenge. In marked contrast to spleen, draining lymph nodes are devoid of transitional, marginal zone and ABC memory cells, and are enriched in cells that underwent extensive class-switching towards IgG subclasses. Furthermore, long-lived germinal center reactions are extensively observed in mediastinal lymph nodes but less frequently in the spleen. This phenomenon seems to be a general feature of acute viral infections as robust germinal center reactions are found 10 weeks after influenza, SARS-CoV-2 and vesicular stomatitis virus infection (Bachmann et al., 1996). The differences between spleen and draining lymph nodes concerning tissue-specific MBC subsets, IgM vs IgG-expressing cells, and long-lived germinal center reactions add an extra layer of complexity when analyzing recall responses. Future studies investigating the potential of MBCs to re-enter germinal center reactions or acquire a plasma cell program should take these parameters into consideration at the time of choosing the antigen model, the route of immunization and the organ of analysis.

The presence of MBCs is not restricted to secondary lymphoid organs. In recent years, independent studies have shown that MBCs can further settle in non-lymphoid tissues upon resolution of the primary immune response (Oh et al., 2019; Trivedi et al., 2019). For instance, respiratory infection with influenza virus leads to the development of lung-resident MBCs, which persist in the lung mucosa for long periods of time (Allie et al., 2019; Joo et al., 2008; Onodera et al., 2012). We found that these lung-resident MBCs are not uniformly distributed across the lung parenchyma. Instead, MBCs form clusters in close proximity to lung bronchi, a strategic place that allows the rapid encounter of pathogens upon re-challenge and that ensures a first layer of protection directly at the tissue barrier. The positioning of lung MBCs directly at sites of antigen entry resembles the one observed in lymphoid organs, where they preferentially accumulate in lymph node interfollicular/subcapsular sinus areas and splenic marginal/follicular zones (Elgueta et al., 2015; Moran et al., 2018). To be noted, the Ccr6 and Cxcr3 axes are, per se, dispensable for the recruitment/maintenance of MBCs in the lung parenchyma and lymphoid organs (Suan et al., 2017). Then it is plausible that other chemokine pathways, such as the Cxcr5-Cxcl13 axis, are driving this process (Denton et al., 2019). Interestingly, lung-resident MBCs seem to be more cross-protective than their lymphoid tissue counterparts (Adachi et al., 2015; Tamura et al., 1992). While the basis for this phenomenon remains to be determined, this physiological advantage underscores the potential of harnessing B cells directly at the barrier surfaces to generate broadly protective vaccines.

The aforementioned studies, based the identification of lung-resident MBCs on their ability to bind fluorescently-labelled influenza proteins. Here, we took an unbiased fate mapping approach in mice to track MBCs independently of their antigen specificity and based on their early expression of Aid. By doing so, we were able to identify MBCs with a wide range of antigen specificities, including those with no specificity for viral antigens that were disregarded in previous studies. Through unsupervised scRNA-seq analysis we found that the MBC pool in lungs and draining lymph nodes resemble each other at the level of cluster composition and BCR sequences, indicating that these two compartments are linked in their origin. Furthermore, this analysis unveiled the presence of three discrete clusters of MBCs across lungs and lymph nodes, each of them representing a MBC subset with unique expression patterns of Ccr6 and Cxcr3 receptors. The smallest subset consists of innate-like MBCs, which lack Ccr6 and Cxcr3 expression and exclusively express IgM. We found that these cells are localized at the periphery of lung peribronchial B cell clusters, do not show apparent specificity for viral antigens, and resemble splenic B1-like MBCs based on their gene expression and BCR sequences. These cells may arise from spontaneous germinal center reactions in Peyer’s patches or spleen, as a recent study has shown that a large population of IgM^+^ MBCs bearing B1 cell markers, a BCR repertoire skewed towards IgHV6 and specificity for commensal bacteria emerge from there (Le Gallou et al., 2018). Besides this minor subset, two large subsets of Ccr6-expressing MBCs further emerge after influenza infection, one of them lacking Cxcr3 expression and the other one characterized by high Cxcr3 surface levels.

We found that both Cxcr3^-^ and Cxcr3^+^ MBC subsets undergo extensive class-switching towards IgG subclasses, share the peribronchial niche in the lung mucosa, and arise from germinal centers, based on their CD40-L dependence and somatic hypermutation levels. These subsets preferentially differentiate into plasma cells, rather than to germinal center cells, and produce similar levels of IgG when challenged ex vivo. Thus, these two transcriptionally distinct memory populations do not segregate cells regarding fate but instead, cells generated through divergent mechanisms. In fact, the Cxcr3^+^ subset constitutes *bona fide* MBCs, which are mostly pathogen-specific and are actively recruited during secondary immune responses, while the Cxcr3^-^ subset represent *bystander* MBCs with no evident specificity for pathogen-derived antigens. The following questions are inexorable: how are MBCs from the Cxcr3^-^ subset selected in the germinal center and what are they specific for? Elegant work by Kelsoe and collaborators showed that clonal selection is highly permissive in response to complex antigens, allowing the presence of B cells within germinal center reactions with a broad range of affinities and even cells with no detectable affinity for native immunogens (Kuraoka et al., 2016). While the authors showed that these “less-fit” germinal center B cells showed extensive somatic hypermutation and clonal diversification, it remained unknown whether they could acquire a memory phenotype. Here, we show that these cells not only enter the long-lived memory compartment but also that they express a unique transcriptional program that distinguishes them from *bona fide* MBCs. Regarding antigen specificity, it seems unlikely that these cells are very-low affinity, as they acquired VH mutations at comparable if not higher rates than bona fide MBCs and do not display clonal overlap with antigen-specific cells. A potential explanation could be that these cells recognize common autoantigens released as a consequence of immunization. Still, the abovementioned “less-fit” germinal center cells induced by complex antigens do not bind to autoimmune nor common microbial antigens (Horns et al., 2020; Kuraoka et al., 2016). An alternative could be that these cells recognize non-native conformations of immunogens, such as cryptic epitopes exposed by degradation or neoepitopes generated by complement fixation. Yet, our in vivo data showing that Cxcr3^-^ MBCs do not significantly engage into secondary influenza responses, do not go in line with this hypothesis.

Then, how do these cells still make it to the memory compartment? It has been proposed that low-affinity B cells remain in germinal center reactions due to bystander help received from adjacent T follicular helper cells activated by fitter B cells (Wan et al., 2019; Yeh et al., 2018; Zaretsky et al., 2017). Furthermore, recent work has shown that low affinity cells express high levels of CD40 on the surface, presumably to compensate for insufficient T cell help (Nakagawa et al., 2021). While these premises could explain the maintenance of low-affinity B cells in germinal center reactions, it seems unlikely that these processes would keep B cells with no apparent specificity for the immunogen. An alternative explanation arises from our RNA-seq data, which revealed that the Cxcr3^-^ MBC subset expresses high levels of *Fcer2a* and *Fcmr*, the Fc receptors for IgE and IgM. It is tempting to speculate that certain non-specific B cells, expressing unusually high levels of Fc receptors, can internalize immune complexes, present antigen on their surface, and recruit T cell-help independently of their original antigen specificity. This phenomenon, which has been well characterized in the context of allergic responses, could offer an explanation to why *bona fide and bystander* MBCs display divergent transcriptional programs, linked to the presence or absence of BCR signalling (Villazala-Merino et al., 2020). Extensive work is required to delineate the role of Fc receptors on germinal center selection and acquisition of divergent MBC programs.

Permissive selection seems to be a general strategy of the immune system activated upon pathogen encounter, as we found that SARS-CoV-2 infection also engenders *bona fide* and *bystander* MBC subsets with contrasting antigen specificities. Regardless of the mechanism driving the formation of these memory subsets, our results raise the important question of the potential benefit of such phenomenon, as *bystander* cells can represent up to 50% of the MBC pool generated upon infection. We propose that the presence of high and low-affinity MBCs within the *bona fide* subset helps the host to overcome future infections with the same pathogen, mutating variants or even different viruses from the same family, as recently observed with flaviviruses (Viant et al., 2020; Wong et al., 2020). In contrast, the prevalence of *bystander* MBCs with no detectable specificity for the infecting pathogen could be beneficial to expand the diversity of the initial B cell repertoire while acting as platforms to retain immune complexes in close proximity to *bona fide* MBCs. We speculate that this phenomenon could operate as a safeguard mechanism to counteract the loss in B cell repertoire diversity observed, for instance, upon ageing (Gibson et al., 2009; Siegrist and Aspinall, 2009).

## Materials and methods

### Mice

8-week old wild-type C57BL/6 mice were obtained from Janvier Labs. Aicda-Cre^ERT2^ mice were obtained from Claude-Agnès Reynaud and Jean Claude Weill, Institut Necker Enfants Malades, France. Rosa26-EYFP, CCR6^-/-^ and K18-hACE2 mice were obtained from Jackson Laboratories, USA. μMT mice were obtained from Stéphane Mancini, Centre de Recherche en Cancérologie de Marseille, France. CXCR3^-/-^ and Fcmr^-/-^ bone marrow were obtained from Jacqueline Marvel, Centre International de Recherche en Infectiologie, France and Tak Mak, University of Toronto, Canada, respectively. Aicda-Cre^ERT2^ mice were further crossed with Rosa26-EYFP and CCR6^-/-^ mice.

For the generation of mixed bone marrow chimeras, μMT mice of 6-8 weeks of age were irradiated with 2 doses of 4.75 Gy, 4 hours apart. One day later, bone marrow cells were injected *i*.*v*. in recipient animals (1×10^6^ cells in total). Experimental animals were kept on water with Bactrim for 3 days prior and 3 weeks post irradiation treatment. Chimeras were used after 8 weeks of reconstitution.

Mice were bred and maintained at the animal facilities of the Centre d’Immunologie de Marseille Luminy and Centre d’Immunophénomique. Up to five mice per cage were housed under a standard 12 hr light/dark cycle, with room temperature at 22°C (19°C-23°C change). They were fed with autoclaved standard pellet chow and reverse osmosis water. All cages contained 5 mm of aspen chip and tissue wipes for bedding and a mouse house for environmental enrichment. Mice were used at the age of 8 to 12 weeks and littermates (males or females) were randomly assigned to experimental groups. Generally between 4 to 8 mice were used per experimental group. Experimental procedures were conducted in accordance with French and European guidelines for animal care under the permission number 16708-2018091116493528 following review and approval by the local animal ethics committee in Marseille.

### Infections and injections

Mice were anesthetized i.p. with Ketamine/Xylazine (100 mg/kg body) and intranasally infected with 5 PFU of Influenza virus A/Puerto Rico/8/1934 (PR8) H1N1 strain or 10^4^ PFU of Influenza virus A/X-31 H3N2 or 10^2^-10^4^ PFU of SARS-CoV-2 in 20 μl of PBS. In indicated experiments, Influenza PR8 was inactivated under 365 nm long-wave UV light for 10 minutes and administered intranasally at 10^5^ PFU/mouse. Influenza virus was amplified on MDCK cells. Purification of viral particles was performed in a sucrose 30% cushion at 25,000RPM for 2 hours in an SW32Ti rotor. For Aid-EYFP mice, tamoxifen (Cayman chemical) was resuspended in corn oil (Sigma), sonicated and given by oral gavage at 5mg/100μl. For in vivo disruption of germinal centers, mice were i.p. injected with 300 μg of anti-CD40L (MR1, BioXCell) in 200 μl of PBS at different days of infection. For in vivo labelling of immune cells in circulation, 3 μg of anti-CD45 antibody was administered i.v 5 minutes before sacrifice. For adoptive transfers, bone marrow cells (1.10^6^) were resuspended in 100 μl of PBS and i.v. injected. For the protection experiment, μMT mice received 5.10^6^ B6 splenocytes i.v. and 15μg of CpG ODN-1826 (Invivogen) intranasally in PBS (Onodera et al., 2012). After 10 days, mice received 10^6^ splenic CD4^+^ T cells and 3.10^3^ Ccr6^+^Cxcr3^-^ or Ccr6^+^Cxcr3^+^ MBCs sorted from lungs of B6 mice infected with Influenza PR8 70 days before. The day after, mice were challenged with PR8 virus, weight was measured daily and euthanized if exhibiting ≥20% loss of initial mass.

The strain BetaCoV/France/IDF0372/2020 (SARS-CoV-2) was supplied by the National Reference Centre for Respiratory Viruses hosted by Institut Pasteur (Paris, France) and headed by Pr. Sylvie van der Werf. The human sample from which strain BetaCoV/France/IDF0372/2020 was isolated has been provided by Dr. X. Lescure and Pr. Y. Yazdanpanah from the Bichat Hospital, Paris, France. Moreover, the strain BetaCoV/France/IDF0372/2020 was supplied through the European Virus Archive goes Global (Evag) platform, a project that has received funding from the European Union’s Horizon 2020 research and innovation programme under grant agreement No 653316. Infectious stocks were grown by inoculating Vero E6 cells and collecting supernatant upon observation of cytopathic effect; debris were removed by centrifugation and passage through a 0.22-μm filter. Supernatant was then aliquoted and stored at −80 °C. Vero E6 (CRL-1586; American Type Culture Collection) were cultured at 37 °C in Dulbecco’s modified Eagle’s medium (DMEM) supplemented with 10% fetal bovine serum (FBS), 10 mM HEPES (pH 7.3), 1 mM sodium pyruvate, 1 x non-essential amino acids and 100 U ml−1 penicillin–streptomycin. Work with SARS-CoV-2 was performed in the biosafety level 3 laboratory of Center for Immunophenomics (CIPHE) by personnel equipped with powered air-purifying respirators. All the CIPHE BSL3 facility operations are overseen by a Biosecurity/Biosafety Officer and accredited by Agence Nationale de Sécurité du Médicament (ANSM).

### Flow Cytometry

Single cell suspensions of mediastinal lymph nodes and spleens were obtained by pressing organs through a 70μm nylon mesh cell strainer with a plastic plunge in PBS 2%FCS 2mM EDTA. For lungs, single cell suspensions were obtained with the mouse lung dissociation kit (Miltenyi) according to manufacturer instructions. For spleen and lungs, cell suspensions were further incubated with red blood cell lysis buffer for 5 minutes. To block nonspecific antibody binding, cell suspensions were incubated with hybridoma supernatant 2.4G2, diluted 1/5 in PBS 2%FCS 2mM EDTA, for 20 minutes on ice. For labeling of surface markers, cells were stained for 20 minutes on ice with the indicated anti-mouse antibodies and fluorescently-labelled influenza hemagglutinin, nucleoprotein or SARS-CoV-2 spike protein (Sino biological). Labelling of recombinant proteins was carried out using APC and PE conjugation lightning-link kits (Abcam). When using biotinylated antibodies, cell suspensions were washed following antibody labeling and incubated for 20 minutes on ice with labeled streptavidin. Finally, cells were either resuspended in 400μl of PBS 2%FCS 2mM EDTA and analyzed on Fortessa-X20/Symphony cytometers (BD Biosciences) or in RPMI media and used for cell sorting on a FACSAria II (BD, Bioscience). DAPI (0.1 μg/ml final concentration) was added right before passing samples on cytometers. In the case of samples from SARS-CoV-2 infected animals, cell suspensions were incubated with Zombie UV fixable viability kit (Biolegend) to exclude dead cells and samples were fixed 30 minutes with PFA 4%. Data was analyzed using FlowJo (TreeStar).

| Antibody | Source | Catalog |
| --- | --- | --- |
| APC/Cy7 Rat Anti-Mouse CD19, Clone 6D5 | Biolegend | 115530 |
| BUV395 Rat Anti-Mouse CD38, Clone 90/CD38 | BD | 740245 |
| PE/Cy7 Rat Anti-Mouse CD38, Clone 90 | Biolegend | 102718 |
| Alexa Fluor 647 Rat Anti-Mouse/Human GL7 Antigen, Clone GL7 | Biolegend | 144606 |
| PerCP/Cy5.5 Rat Anti-Mouse/Human GL7 Antigen, Clone GL7 | Biolegend | 144610 |
| PE Rat Anti-Mouse CD138 (Syndecan-1), Clone 281-2 | Biolegend | 142504 |
| BV785 Rat Anti-Mouse CD138 (Syndecan-1), Clone 281-2 | Biolegend | 142534 |
| BV650 Rat Anti-Mouse CD93, Clone AA4.1 | BD | 563807 |
| BV605 Mouse Anti-Mouse CD95 (Fas), Clone SA367H8 | Biolegend | 152612 |
| PE/Cy7 Rat Anti-Mouse IgD, Clone 29-2L17 | Biolegend | 405720 |
| BV786 Rat Anti-Mouse IgD, Clone 11-26c.2a | BD | 563618 |
| FITC Rat Anti-Mouse CD45, Clone 30-F11 | Biolegend | 103108 |
| PE Rat Anti-Mouse CD45, Clone 30-F11 | Biolegend | 103106 |
| PE/Dazzle 594 Armenian Hamster Anti-Mouse CD196 (CCR6), Clone 29-2L17 | Biolegend | 129822 |
| BV421 Armenian Hamster Anti-Mouse CD183 (CXCR3), Clone CXCR3-173 | Biolegend | 126522 |
| BV605 Armenian Hamster Anti-Mouse CD11c, Clone N418 | Biolegend | 117334 |
| BV510 Rat Anti-Mouse CD21/CD35, Clone 7G6 | BD | 747764 |
| PE/Cy7 Armenian Hamster Anti-Mouse CD55 (DAF), Clone RIKO-3 | Biolegend | 131814 |
| PE/Cy5 Rat Anti-Mouse CD24, Clone M1/69 | Biolegend | 101812 |
| PerCP/Cy5.5 Rat Anti-Mouse IgM, Clone RMM-1 | Biolegend | 406512 |
| V450 Rat Anti-Mouse IgG1, Clone A85-1 | BD | 562107 |
| PE Rat Anti-Mouse IgG2b, Clone RMG2b-1 | Biolegend | 406708 |
| Biotinylated Goat Anti-Mouse IgG2c, polyclonal | Southern BT | 1079-08 |
| BV650 Rat Anti-Mouse IgG3, Clone R40-82 | BD | 744136 |
| Alexa Fluor 647 Goat Anti-Mouse IgA, polyclonal | Southern BT | 1040-31 |
| PerCP/Cy5.5 Armenian Hamster Anti-Mouse CD3e, clone 145-2C11 | Biolegend | 100328 |
| BV605 Rat Anti-Mouse CD4, clone RM4-5 | Biolegend | 100548 |
| APC/Cy7 Rat Anti-Mouse CD8a, clone 53-6.7 | Biolegend | 100714 |
| PE Rat Anti-Mouse/Human CD44, clone IM7 | Biolegend | 103008 |
| APC Rat Anti-Mouse CD62L, clone MEL-14 | Biolegend | 104412 |
| PE/Cy7 Armenian Hamster Anti-Mouse CD69, clone H1.2F3 | Biolegend | 104512 |
| BV711 Armenian Hamster Anti-Mouse CD103, clone 2E7 | Biolegend | 121435 |
| APC Rat Anti-Mouse CD23, clone B3B4 | Biolegend | 101620 |
| Purified Mouse Anti-Mouse Fcμr, clone MM3 | Biolegend | 157802 |

### Immunohistochemistry

Lungs, lymph nodes and spleens were fixed in 4% PFA for 6 hours at 4°C, washed with PBS, incubated overnight in PBS 30% sucrose solution, immersed in OCT and snap-frozen in liquid nitrogen-cooled isopentane. Cryostat sections (10 to 20 µm thick) were dried in silica beads, permeabilized with PBS Saponin 0.5% for 30 minutes and blocked with PBS 0.5% saponin 2% BSA 1% goat serum 1% FCS for 30 minutes. Sections were then incubated with primary antibodies in PBS 0.5% saponin 2% BSA 1% goat serum 1% FCS for at least 1 hour, washed and incubated with secondary antibodies for a further hour. After a final wash, sections were mounted in Fluoromount-G mounting media. Imaging was carried out on a LSM 780 (Zeiss) inverted confocal microscope using a Plan-Apochromat 40x NA 1.3 oil immersion objective or a Plan-Apochromat 20X/0.8 M27 objective in the case of full organ section.

| Antibody | Source | Catalog |
| --- | --- | --- |
| Pacific Blue Rat Anti-Mouse/Human CD45R (B220), Clone RA3-6B2 | Biolegend | 103227 |
| Alexa Fluor 488 Rat Anti-Mouse CD326 (Ep-CAM), Clone G8.8 | Biolegend | 118210 |
| Alexa Fluor 647 Rat Anti-Mouse CD326 (Ep-CAM), Clone G8.8 | Biolegend | 118212 |
| APC Rat Anti-Mouse CD3, clone 17A2 | Biolegend | 100236 |
| Alexa Fluor 647 Rat Anti-Mouse/Human GL7 Antigen, Clone GL7 | Biolegend | 144606 |
| PE Rat Anti-Mouse CD138 (Syndecan-1), Clone 281-2 | Biolegend | 142504 |
| Alexa Fluor 647 Rat Anti-Mouse CD169 (Siglec-1), Clone 3D6.112 | Biolegend | 142408 |
| PE/Dazzle 594 Armenian Hamster Anti-Mouse CD196 (CCR6), Clone 29-2L17 | Biolegend | 129822 |
| APC Armenian Hamster Anti-Mouse CD183 (CXCR3), Clone CXCR3-173 | Biolegend | 126512 |
| eFluor 570 Rat Anti-Mouse IgM, Clone II/41 | eBioscience | 41579082 |
| PE Rat Anti-Mouse IgG2b, Clone RMG2b-1 | Biolegend | 406708 |
| Alexa Fluor 488 Rat Anti-Mouse IgG2b, Clone RMG2b-1 | Biolegend | 406718 |
| Chicken Anti-GFP, polyclonal | Abcam | ab13970 |
| Alexa Fluor 488 Goat Anti-Chicken IgY (H+L), polyclonal | Invitrogen | A11039 |
| Rabbit Anti-SARS-CoV-2 Nucleocapsid, polyclonal | GeneTex | GTX135357 |
| Alexa Fluor 555 Goat Anti-Rabbit IgG (H+L), | Invitrogen | A21428 |

### Analysis of confocal images

Image segmentation was performed using a specific ImageJ macro based on classical binary watershed to generate a mask on YFP^+^ cells. Selections were corrected manually by overlay masking. Mean fluorescence intensity for the B220, CXCR3 and CCR6 channels as well as the area in each COI (Cell of Interest) was exported. The ROI (Region of interest) was defined manually at the beginning of the macro and exported in order to calculate cell density. The macro exports 4 files: the image in TIF, the COI in zip format (.roi), the mean fluorescence intensity and area for each cell in csv, and the channel legend. A specific software named SAPHIR (Shiny Analytical Plot of Histological Image Results) was developed using R, the code source can be downloaded here: https://github.com/elodiegermani/SAPHIR. This software uses the 4 files exported by the ImageJ macro and allows scatterplot gating (cross, polygon or lasso) to filter and define cell populations interactively with the COI overlaid on the image.

### Ex vivo culture of MBCs on 40LB cells

40LB cells were cultured in DMEM supplemented with 10% FCS, penicillin 100u/ml, streptomycin 100µg/ml, sodium pyruvate 1mM and beta-mercaptoethanol 5×10^−5^M as previously described (Nojima et al., 2011). 40LB cells were irradiated (120Gy) and seeded in 24-well plates at a density of 0.25×10^6^ cells/well. The following day, sorted MBCs were added on irradiated feeder cells in RPMI, 10% FCS, penicillin 100u/ml, streptomycin 100µg/ml, sodium pyruvate 1mM and beta-mercaptoethanol 5×10^−5^M. Mouse recombinant IL-21 (PeproTech) was added to co-cultures at 10ng/ml. At indicated time points, cells were analyzed by flow cytometry and supernatant collected for posterior antibody analysis.

### ELISPOT

To measure influenza-specific antibody-secreting cells, enzyme-linked immunosorbent spot (ELISPOT) multiscreen filtration plates (Millipore) were activated with absolute ethanol, washed with PBS and coated overnight at 4°C with 1μg/ml of nucleoprotein (Sino Biological) diluted in PBS. Plates were subsequently blocked for 1 hour with complete medium and incubated for 24 hours at 37°C with serial dilutions of lymph node, lung and spleen single cell suspensions. For lungs, the lymphocyte fraction was previously enriched using 40:80% Percoll (GE Healthcare). Plates were washed with PBS 0.01% Tween-20 and incubated for 1 hour with 1μg/ml of biotinylated anti-IgM, IgG or IgA (Southern Biotech) diluted in PBS 1% BSA. Then, plates were washed and incubated for 30 minutes with 1μg/ml Streptavidin-Alkaline Phosphatase (Sigma). Finally plates were washed and developed with BCIP®/NBT (Sigma).

### ELISA

To measure influenza-specific antibodies, ELISA (enzyme-linked immunosorbent assay) plates were coated overnight at 4°C with 1μg/ml of nucleoprotein or hemagglutinin (Sino Biological) diluted in PBS. Plates were washed with PBS 0.01% Tween, blocked for 2 hours at room temperature with PBS 2.5% FCS and incubated overnight at 4°C with serial dilutions of co-culture supernatants. The following day, plates were washed and probed at room temperature for 1 hour with 1μg/ml of biotinylated anti-IgG (Southern Biotech) in PBS 2.5%FCS. Plates were washed and incubated at room temperature for 30 minutes with 1μg/ml Streptavidin-Alkaline Phosphatase (Sigma). Plates were washed and developed with p-NitrophenylPhosphate (Sigma). 405nm absorbance was detected using a SPECTRAmax190 plate reader (Molecular Device).

### 10x 5’ scRNA-Seq library preparation

In a first scRNA-seq experiment (Figures 2 and 4), single-cell suspensions from spleen, lymph nodes and lungs were prepared as described above. Because enzymatic digestion of tissues at 37°C induced the expression of dissociation-induced genes in some lung MBCs (Extended Figure 2), we performed a second experiment using only mechanical dissociation of lung tissue at 4°C. For both experiments, we used cell hashing with hashtag oligonucleotides (HTO) to multiplex 3 organ samples from 3 individual mice as previously described (Mimitou et al., 2019). After cell surface staining with the mix of antibodies used for gating MBCs, single-cell suspensions from each organ of each individual mice were independently stained with a distinct barcoded anti-mouse CD45 antibody (in-house conjugated) in PBS 2%FCS 2mM EDTA for 30 min on ice, then washed and resuspended in PBS. For each sample, live MBCs (DAPI^-^CD19^+^eYFP^+^CD38^+^GL7^-^) were bulk-sorted with BD FACS Aria II. Sorted cell samples (experiment 1: 13,126 lung cells, 3,414 lymph node cells, 12,838 spleen cells; experiment 2: 1,319 lung cells, 1,436 lymph node cells, 1,500 spleen cells) were pooled and loaded in a single capture well for subsequent 10x Genomics Single Cell 5’ v1 workflow.

10x 5’ scRNA-seq libraries were prepared according to the manufacturer’s instructions with modifications for generating the BCR-seq libraries. Following cDNA amplification, SPRI select beads were used to separate the large cDNA fraction derived from cellular mRNAs (retained on beads) from the HTO-containing fraction (in supernatant). For the cDNA fraction derived from mRNAs, 50ng were used to generate transcriptome library and around 5ng were used for BCR library construction. Gene expression libraries were prepared according to manufacturer’s instructions. For BCR libraries, heavy and light chain cDNA were amplified by two rounds of PCR (6 cycles + 8 cycles) using external primers recommended by 10x Genomics, and 800 pg of purified amplified cDNA was tagmented using Nextera XT DNA sample Preparation kit (Illumina) and amplified for 12 cycles using the SI-PCR forward primer (10x Genomics) and a Nextera i7 reverse primer (Illumina). For the HTO-containing fraction, 5ng were used to generate the HTO library. The resulting libraries were pooled and sequenced together on an Illumina NextSeq550 platform, using High Output 75-cycle flow cells, targeting 5×10^4^ reads per cell for gene expression, 5×10^3^ reads per cell for BCR, 2×10^3^ reads per cell for hashtag, in paired-end single-index mode (Read 1: 26 cycles, Read i7: 8 cycles, Read 2: 57 cycles).

### FB5P-seq library preparation

Single-cell suspensions from lungs of three previously infected Aid-EYFP mice were prepared as described above with enzymatic digestion, and stained with a panel of antibodies for identifying subsets of antigen-specific MBCs (GL7-PerCP-Cy5.5, HA-PE, Ccr6-PE-Dazzle594, CD38-PE-Cy7, NP-APC, CD19-APC-Cy7, Cxcr3-BV421, Live/Dead Aqua stain). FB5P-seq protocol was performed as previously described (Attaf et al., 2020). Briefly, single MBCs were FACS sorted on a BD Influx into 96-well PCR plates containing 2µl lysis mix per well. The index-sorting mode was activated to record the different fluorescence intensities of each sorted cell. Immediately after cell sorting, plates containing single cells in lysis mix were frozen on dry ice and stored at −80°C until further processing. For each plate, library preparation consisted in RT with template switching for incorporating well-specific barcodes and UMIs, cDNA amplification with 22 cycles of PCR, pooling of 96 wells into one tube, and 5’-end RNA-seq library preparation using tagmentation-based modified Nextera XT DNA sample Preparation kit (Illumina). Libraries were tagged with a plate-specific i7 index and were pooled for sequencing on an Illumina NextSeq2000 platform, with P2 flow cells, targeting 2.5×10^5^ reads per cell in paired-end single-index mode (Read 1: 103 cycles, Read i7: 8 cycles, Read 2: 16 cycles).

### scRNA-seq analysis

Preprocessing and analysis of data were done through the usage of standard tools and custom R and *Python* scripts. We notably used R version 3.5 and 3.6, Seurat package version 3 (Stuart et al., 2019), 10x Genomics CellRanger version 3, CITE-seq-count version 1.4 (Stoeckius et al., 2018). Docker and Singularity containers were used to ensure the reproducibility of analyses. All codes and data are available on Github and Zenodo.

#### Pre-processing of FB5P-seq dataset

We used a custom bioinformatics pipeline to process fastq files and generate single-cell gene expression matrices and BCR sequence files as previously described (Attaf et al., 2020). Detailed instructions for running the FB5P-seq bioinformatics pipeline can be found at https://github.com/MilpiedLab/FB5P-seq. Quality control was performed to remove poor quality cells. Cells with less than 500 genes detected and genes detected in at least 3 cells were removed. We further excluded bad quality cells expressing more than 3% of mitochondrial genes or less than 10% of ribosomal genes. For each cell, gene expression UMI count values were log-normalized with Seurat *NormalizeData* with a scale factor of 10,000 (Stuart et al., 2019).

Index-sorting FCS files were visualized in FlowJo software and compensated parameters values were exported in CSV tables for further processing. For visualization on linear scales in the R programming software, we applied the hyperbolic arcsine transformation on fluorescence parameters (Finak et al., 2010).

For BCR sequence reconstruction, the FB5P-seq pipeline used Trinity for *de novo* transcriptome assembly for each cell based on Read1 sequences, then MigMap for filtering the resulting contigs for productive BCR sequences and identifying germline V, D and J genes and CDR3 sequence for each contig. Filtered contigs were aligned to reference constant region sequences using Blastn. The FB5P-seq pipeline also ran the pseudoaligner Kallisto to map each cell’s FB5P-seq Read1 sequences on its reconstructed contigs and quantify contig expression. The outputs of the FB5P-seq pipeline were further processed and filtered with custom R scripts. For each cell, reconstructed contigs corresponding to the same V(D)J rearrangement were merged, keeping the largest sequence for further analysis. We discarded contigs with no constant region identified in-frame with the V(D)J rearrangement. In cases where several contigs corresponding to the same BCR chain had passed the above filters, we retained the contig with the highest expression level. BCR metadata from the MigMap and Blastn annotations were appended to the gene expression and index sorting metadata for each cell. Finally, heavy and light chain contig sequences were trimmed to retain only sequences from FR1 to the first 36 nucleotides of FR4 regions, and were exported as fasta files for further analysis of clonotypes and BCR phylogenies.

#### Pre-processing of 10x 5’ datasets

Raw fastq files from gene expression libraries were processed using Cell Ranger software, with alignment on the mm10 reference genome. For each experiment, cells with less than 200 genes detected and genes detected in less than 3 cells were removed. We further excluded bad quality cells expressing less than 2,000 UMI, more than 40% mitochondrial genes or less than 10% ribosomal genes. HTO barcodes for sample demultiplexing after hashing were counted using CITE-seq-count and were normalized for each cell using a centered log ratio (CLR) transformation across cells implemented in the Seurat function *NormalizeData*. Cells were demultiplexed using Seurat *MULTIseqDemux* function and barcodes assigned as doublets or negative were excluded from further analysis. The resulting filtered UMI count matrices were log-normalized with Seurat *NormalizeData* with a scale factor of 10,000.

BCR-seq raw fastq files were processed with the FB5P-seq pipeline (Attaf et al., 2020) as described above for FB5P-seq datasets, omitting the part of the pipeline related to gene expression analysis, and using the list of cell-associated 10x barcodes from CellRanger analysis as inputs for splitting bam files upstream Trinity assembly of BCR contigs. BCR metadata from the MigMap and Blastn annotations were appended to the gene expression metadata for each cell. Heavy and light chain variable sequences, trimmed to retain only sequences from FR1 to the first 36 nucleotides of FR4 regions, were exported as fasta files for further analysis of clonotypes and BCR phylogenies.

#### BCR-seq based phylogenies

For selected large clones in the FB5P-seq dataset, we inferred the unmutated common ancestor (UCA) sequence by combining the IMGT-defined germline V_L_, J_L_ and V_H_, D_H_, J_H_ sequences with the observed V_L_-J_L_ and V_H_-D_H_-J_H_ junctional sequences from the least somatically mutated sequence observed in the clonotype. For phylogenetic analyses, concatenated variable (FR1 to FR4) IGH and IGL sequences of UCA and all clonally related FL cells were trimmed to equal length and aligned with the GCtree software (DeWitt et al., 2018). Phenotypic group metadata were used as labels to color the different nodes and leaves of the resulting BCR sequence phylogenetic trees.

#### Dataset analysis

Analysis of datasets were performed using custom R scripts. Variable genes (n=2000) were identified with Seurat *FindVariableFeatures (vst* method), BCR coding genes were excluded from the lists of variable genes. After centering with Seurat *ScaleData*, principal component analysis was performed on variable genes with Seurat *RunPCA*, and embedded in two-dimensional UMAP plots with Seurat *RunUMAP* on 20 principal components. UMAP embeddings colored by sample metadata or clusters were generated by Seurat *DimPlot*, those colored by single gene expression or module scores were generated by Seurat *FeaturePlot*, those colored by BCR sequence metadata were generated with ggplot2 *ggplot*.

Clustering was performed using the *FindNeighbors* and *FindClusters* methods of the Seurat package using 30 PC for SNN graph build and Leiden clustering method with sensitivity set to 0.5, 1.3, and 0.5 for the first and second 10x datasets, and the FB5P-seq dataset, respectively.

Marker genes between clusters were identified using the *FindAllMarkers* method of the Seurat package using the Wilcoxon Rank Sum test on genes expressed at least in 10% of the cells, a logFC threshold of 0.25 and a FDR threshold of 0.001.

Heatmap of gene expression along tissue clusters was done performing a mean of the expression of the genes of interest over the clusters, using the *pheatmap* package (version 1.0.12) for the plot. Heatmap of Szymkiewicz–Simpson coefficient were computed using the set of identified marker genes of each cluster and the ggplot2 *ggplot* function (version 3.3) for the plot. Szymkiewicz–Simpson coefficient of an intersection of two sets is the ratio between the size of the intersection between the two sets and the size of the smaller set of the two.

#### Clonotype analysis

Clonotypes were defined using the composition of (i) associated V gene full name (ii) associated J gene full name (e.g. for heavy chain IGHV9-3*01/IGHJ4*01/42 or for light chain IGKV5-48*01/IGKJ2*01/33) of both heavy and light chain (e.g. Clonotype 1 = IGHV9-3*01/IGHJ4*01/42<<>>IGKV5-48*01/IGKJ2*01/33). We used Clustal Omega from *msa* R package (10.1093/bioinformatics/btv494) to evaluate the sequence proximity of clonotypes and verify if full BCR sequences in clonotype were consistent. We observed that distances between sequences within clonotypes were much smaller than distances between clonotypes.

Only cell barcodes found in the transcriptomic data and associated with both light chains and heavy chain information were kept for further analysis. Analysis on shared V genes took into account only the main version of the genes (eg. *IGHV9-3*01* was identified to *IGHV9-3*). Clustal Omega (REF DOI: 10.1093/nar/gkz268) was used to assess the clonotype sequence similarities.

The clonotype overlap heatmap was produced considering the set of clonotypes in each cluster and computing the Szymkiewicz–Simpson coefficient between sets. Hierarchical Clustering was performed using the stats *hclust* function (version 3.6.3) with euclidean distance and average linkage. Plot was performed with the ggplot2 *ggplot* function (version 3.3).

#### Phenotype analysis

Index-sorting flow cytometry data were converted through hyperbolic arcsine transformation (asinh) on fluorescence parameters. A scale factor was applied to each transformation. This scale factor was 20, 10, 100 and 100 for CXCR3, CCR6, HA and NP parameters, respectively. CXCR3^+^, CCR6^+^, HA^+^ and NP^+^ subsets were defined according to the following respective threshold values on asinh transformed data : 0.5, 1.5, 0.8, 0.5.

## Author contributions

C.G. designed and performed experiments, and analyzed data. L.S. analyzed scRNA-seq data. S.V.M. performed in vitro virus assays and flow cytometry analysis. L.G. prepared scRNA-seq libraries. M.M. performed confocal imaging of tissue sections. C.D. performed FB5P-seq and BCRseq data processing. J.M.N. prepared hashtag antibodies. M.F. analyzed confocal images. A.Z. and B.M. provided the K18-hACE2 mouse model and performed SARS-CoV-2 infections. P.M. conceived the project, supervised the scRNA-seq experiments and data analysis. M.G. conceived the project, supervised the work and wrote the manuscript. All authors were involved in scientific discussions.

## Acknowledgements

We thank the flow cytometry, imaging (Imagimm), bioinformatics and genomics platforms for technical support and the biological resource unit for the breeding of animals (CIML). We thank Claude-Agnès Reynaud and Jean Claude Weill for providing Aicda-Cre^ERT2^ mice. We thank Stephane Mancini for providing μMT mice. We thank Jacqueline Marvel for providing Cxcr3^-/-^ bone marrows. We thank Tak Mak for providing Fcmr^-/-^ bone marrows. We thank Ronan Le Goffic for providing influenza virus strains. We thank Daisuke Kitamura and Michel Cogné for sharing 40LB cells and respective protocols. We thank Amandine Sansoni, Philippe Hoest, Lena Gelard, Quentin Bardin, and Marie Malissen for performing and facilitating the studies with SARS-CoV-2 in the CIPHE biosafety level 3. We thank Cathleen Lutz and The Jackson Laboratory for providing K18-hACE2 mice. We thank Sylvie van der Werf for the BetaCoV/France/IDF0372/2020 strain and Bernard La Scola for advice on SARS-CoV2 handling. We thank Sophie Brustlein, Marc Bajenoff and Serge van de Pavert for advice on imaging. We thank Facundo Batista and Pierre Golstein for critical reading of the manuscript and scientific discussions. We acknowledge Centre de Calcul Intensif d’Aix-Marseille for granting access to its high performance computing resources. This work was supported by the ATIP-AVENIR young group leader program (CNRS-INSERM), junior researcher award (INSERM), and Marie Sklodowska Curie reintegration fellowship (EU) to M.G.; Fondation pour la Recherche Médicale fellowship to S.V.M.; ANR-17-CE15-0009-01 (MoDEx-GC) to P.M.; COVIDHUMICE project (Fondation pour la Recherche Médicale-ANR Flash Covid-COVI-0066) to B.M.; ANR-10-INBS-04-01 France Bio Imaging; Centre d’Immunologie de Marseille Luminy (CIML), which receives its core funding from Aix Marseille University, CNRS and INSERM.

## Figure legends

**Extended Figure 1.**
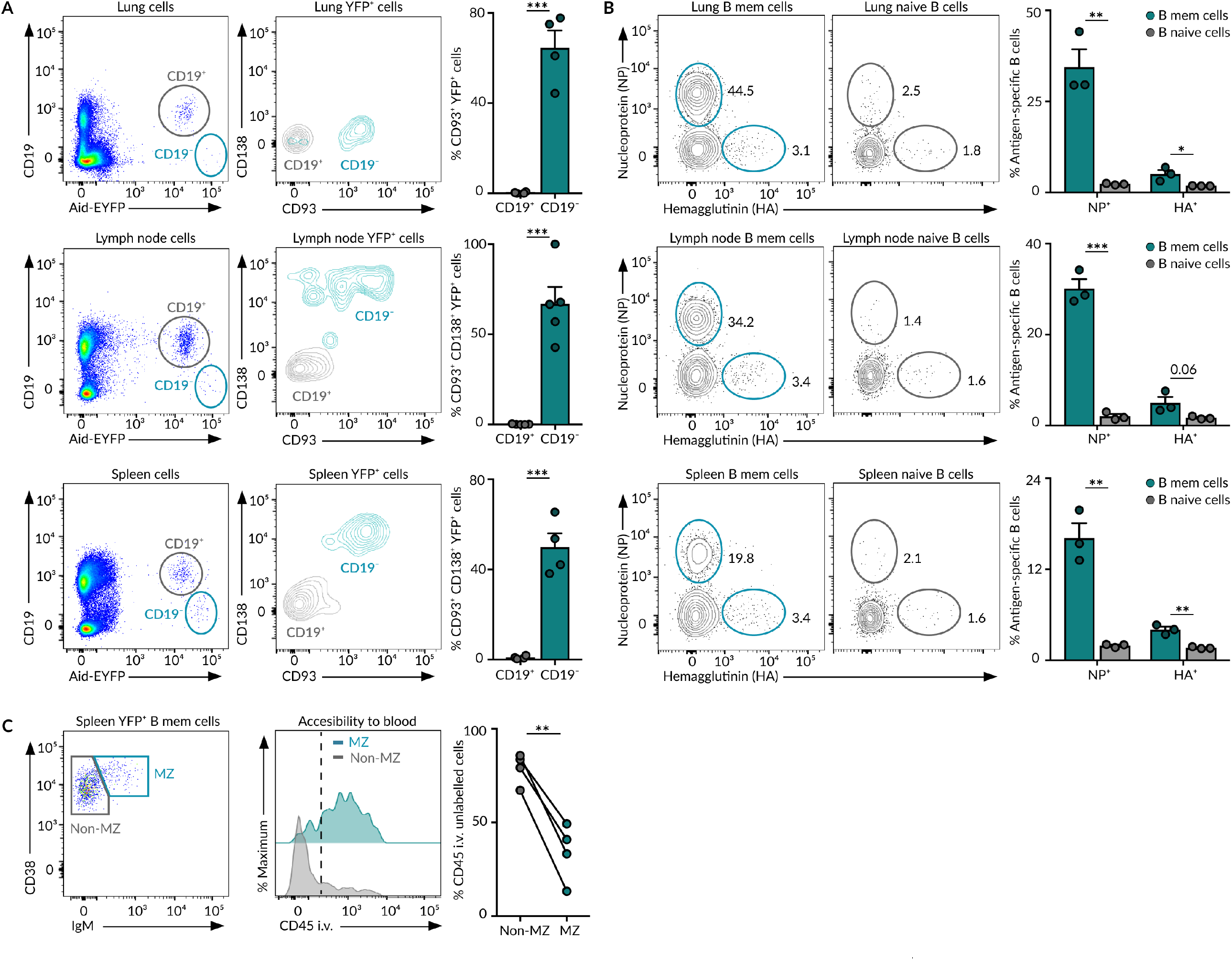
MBCs in influenza infection. **(A)** Contour plots displaying the expression of CD138 and CD93 plasma cell markers by YFP+CD19+ and YFP+CD19-cells in lungs, mediastinal lymph nodes and spleen of Aid-EYFP animals treated as in Figure 1A. Our lung dissociation protocol chops the CD138 epitope. **(B)** Flow cytometry plots showing the percentage of YFP+ MBCs and YFP-IgD+ naive B cells that bind to fluorescently labelled Nucleoprotein (NP) and Hemagglutinin (HA). **(C)** Histogram showing the extent of labelling of YFP+ spleen MBCs, IgM+ and IgM-, with anti-CD45 antibody administered intravenously 5 minutes before sacrifice. Quantification shows the percentage of B cells that are protected from in vivo staining. In panels A and B, bar charts show the quantification of one representative experiment out of three, mean ± s.e.m. Each dot represents one mouse. t test: *p<0.05, **p<0.01 and ***p<0.001.

**Extended Figure 2.**
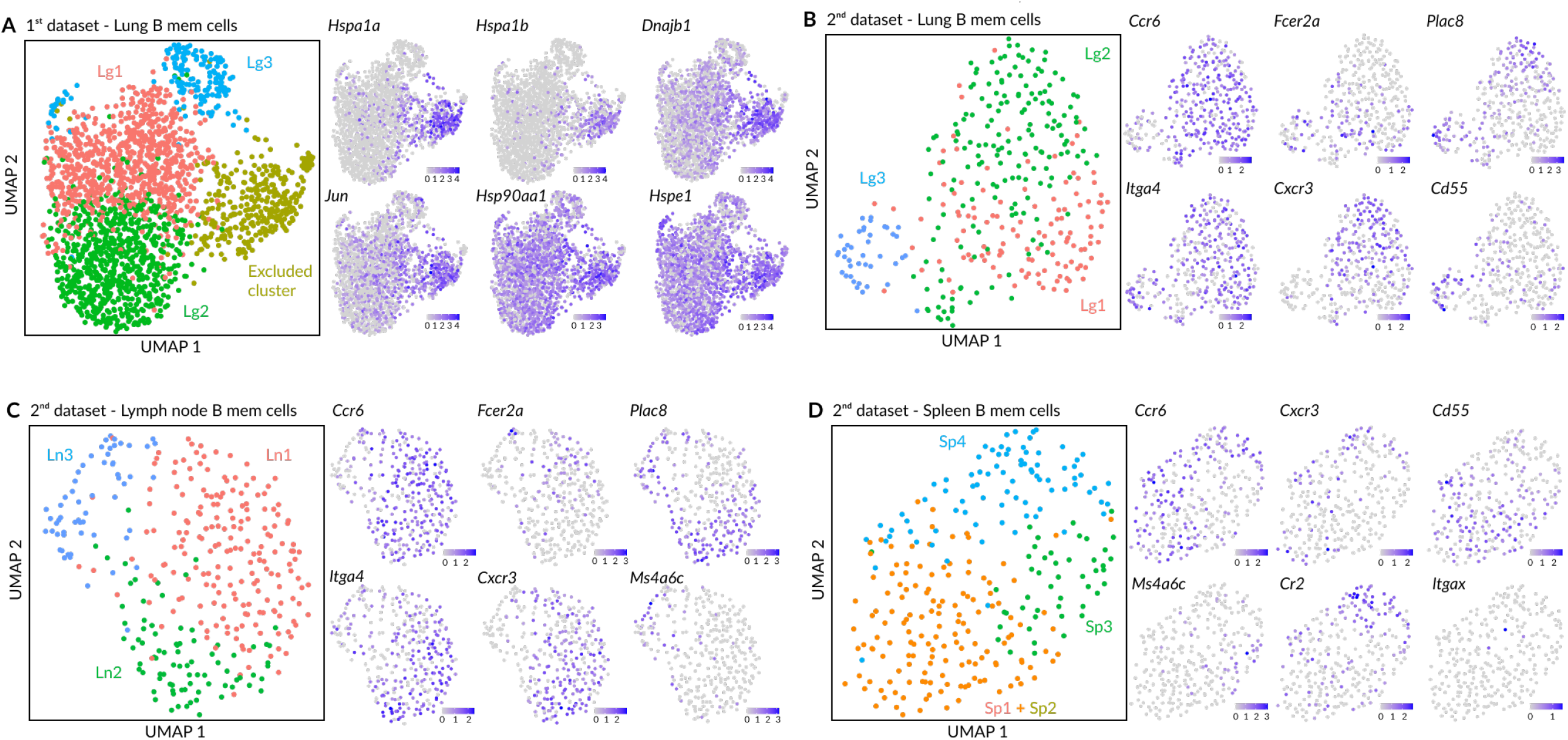
Heterogeneity of MBCs. **(A)** UMAP projection of lung MBCs from the 1^st^ dataset before excluding cells from the cluster with high expression of early-activation genes. In this dataset, lung single cell suspensions were obtained through enzymatic digestion of tissue. **(B-D)** UMAP projections of MBCs in lungs (B), mediastinal lymph node (C) and spleen (D) from the 2^nd^ dataset. In this dataset, lung single cell suspensions were obtained through mechanical disruption of tissue. Due to low cell numbers, spleen MBCs from Sp1 and Sp2 subpopulations clustered together and cells from the Sp5 cluster dispersed across the three clusters identified. In all panels, feature plots display the expression of indicated marker genes laid out in the UMAP representation. Scale: normalized UMI counts.

**Extended Figure 6.**
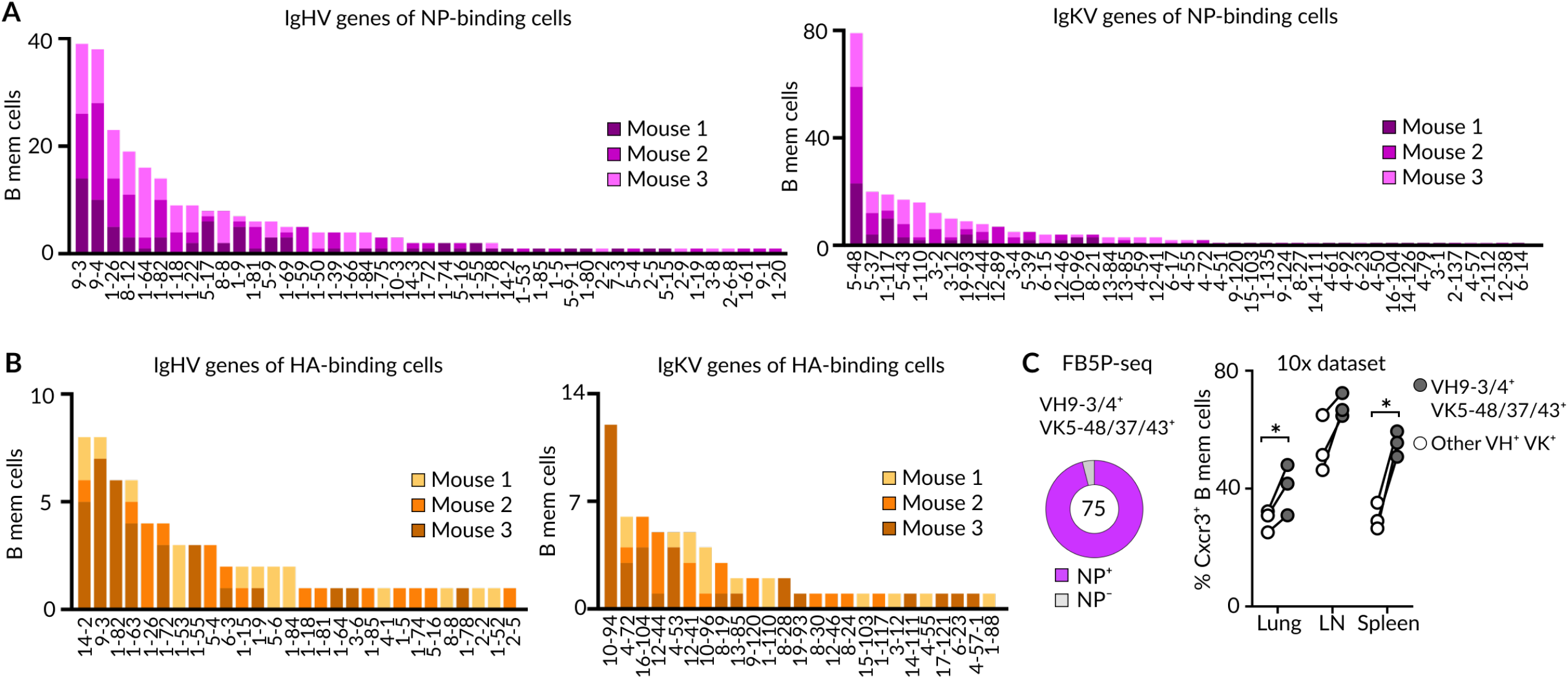
Bona fide and bystander MBCs. **(A-B)** Quantification of IgH and IgK variable gene frequency in MBCs that bind NP (H) or HA (I) from the FB5P-seq data shown in Figure 6. **(C)** Pie chart (left) displaying the percentage of VH9-3/4^+^ VK5-48/37/43^+^ cells that bind NP. Quantification (right) of lung, lymph node and spleen MBCs from the first 10x experiment that show detectable levels of *Cxcr3* transcripts. Cells that bear VH9-3/4^+^ VK5-48/37/43^+^ BCR were compared to cells bearing other BCR combinations, ratio-paired t test, *p<0.05.

**Extended Figure 7.**
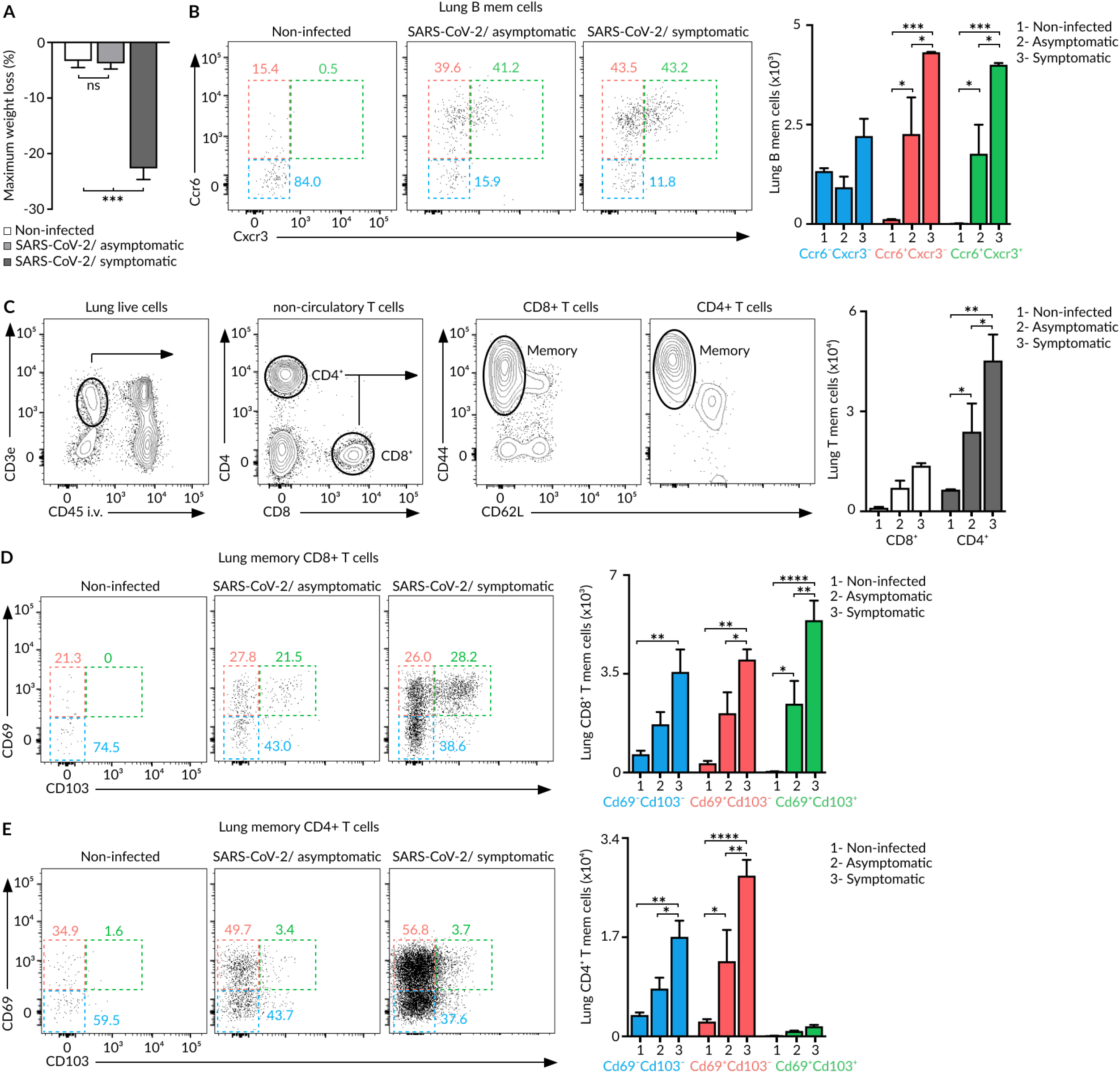
MBC subsets in SARS-CoV-2 infection. **(A)** Bar charts showing the maximum percentage of body weight loss in K18-hACE2 mice that were anaesthetised with ketamine/xylazine and intranasally given PBS (non-infected), 10^1^-10^2^ PFU of SARS-CoV-2 (asymptomatic) or 10^3^-10^4^ PFU of SARS-CoV-2 (symptomatic). **(B)** Dot plots showing the presence of Ccr6^-^ Cxcr3^-^, Ccr6^+^ Cxcr3^-^ and Ccr6^+^ Cxcr3^+^ MBC subsets in lungs of K18-hACE2 mice treated as in (A). **(C)** Flow cytometry plots showing the gating strategy for CD8^+^ and CD4^+^ memory T cells (CD3^+^CD45 i.v.^-^ CD44^+^ CD62L^-^) in lungs of mice treated as in (A). **(D-E)** Dot plots showing the presence of Cd69^-^Cd103^-^, Cd69^+^Cd103^-^ and Cd69^+^Cd103^+^ memory CD8^+^ (D) or CD4^+^ (E) T cell subsets in lungs of K18-hACE2 mice treated as in (A). Bar charts show the quantification of one representative experiment out of three, mean ± s.e.m. Two-way Anova test, *p<0.05, **p<0.01, ***p<0.001 and ****p<0.0001.

## References

Adachi, Y., Onodera, T., Yamada, Y., Daio, R., Tsuiji, M., Inoue, T., Kobayashi, K., Kurosaki, T., Ato, M., and Takahashi, Y. (2015). Distinct germinal center selection at local sites shapes memory B cell response to viral escape. Journal of Experimental Medicine 212, 1709–1723.

Adam, M., Potter, A.S., and Potter, S.S. (2017). Psychrophilic proteases dramatically reduce single-cell RNA-seq artifacts: a molecular atlas of kidney development. Development 144, 3625–3632.

Akkaya, M., Kwak, K., and Pierce, S.K. (2020). B cell memory: building two walls of protection against pathogens. Nat. Rev. Immunol. 20, 229–238.

Allie, S.R., Rameeza Allie, S., Bradley, J.E., Mudunuru, U., Schultz, M.D., Graf, B.A., Lund, F.E., and Randall, T.D. (2019). The establishment of resident memory B cells in the lung requires local antigen encounter. Nature Immunology 20, 97–108.

Anderson, S.M., Tomayko, M.M., Ahuja, A., Haberman, A.M., and Shlomchik, M.J. (2007). New markers for murine memory B cells that define mutated and unmutated subsets. J. Exp. Med. 204, 2103–2114.

Attaf, N., Cervera-Marzal, I., Dong, C., Gil, L., Renand, A., Spinelli, L., and Milpied, P. (2020). FB5P-seq: FACS-Based 5-Prime End Single-Cell RNA-seq for Integrative Analysis of Transcriptome and Antigen Receptor Repertoire in B and T Cells. Front. Immunol. 11, 216.

Bachmann, M.F., Odermatt, B., Hengartner, H., and Zinkernagel, R.M. (1996). Induction of long-lived germinal centers associated with persisting antigen after viral infection. J. Exp. Med. 183, 2259–2269.

Berry, C.T., Liu, X., Myles, A., Nandi, S., Chen, Y.H., Hershberg, U., Brodsky, I.E., Cancro, M.P., Lengner, C.J., May, M.J., et al. (2020). BCR-Induced Ca Signals Dynamically Tune Survival, Metabolic Reprogramming, and Proliferation of Naive B Cells. Cell Rep. 31, 107474.

Denton, A.E., Innocentin, S., Carr, E.J., Bradford, B.M., Lafouresse, F., Mabbott, N.A., Mörbe, U., Ludewig, B., Groom, J.R., Good-Jacobson, K.L., et al. (2019). Type I interferon induces CXCL13 to support ectopic germinal center formation. J. Exp. Med. 216, 621–637.

DeWitt, W.S., 3rd, Mesin, L., Victora, G.D., Minin, V.N., and Matsen, F.A., 4th (2018). Using Genotype Abundance to Improve Phylogenetic Inference. Mol. Biol. Evol. 35, 1253–1265.

Dogan, I., Bertocci, B., Vilmont, V., Delbos, F., Mégret, J., Storck, S., Reynaud, C.-A., and Weill, J.-C. (2009). Multiple layers of B cell memory with different effector functions. Nat. Immunol. 10, 1292–1299.

Elgueta, R., Marks, E., Nowak, E., Menezes, S., Benson, M., Raman, V.S., Ortiz, C., O’Connell, S., Hess, H., Lord, G.M., et al. (2015). CCR6-dependent positioning of memory B cells is essential for their ability to mount a recall response to antigen. J. Immunol. 194, 505–513.

Finak, G., Perez, J.-M., Weng, A., and Gottardo, R. (2010). Optimizing transformations for automated, high throughput analysis of flow cytometry data. BMC Bioinformatics 11, 546.

Gaebler, C., Wang, Z., Lorenzi, J.C.C., Muecksch, F., Finkin, S., Tokuyama, M., Cho, A., Jankovic, M., Schaefer-Babajew, D., Oliveira, T.Y., et al. (2021). Evolution of antibody immunity to SARS-CoV-2. Nature 591, 639–644.

Gibson, K.L., Wu, Y.-C., Barnett, Y., Duggan, O., Vaughan, R., Kondeatis, E., Nilsson, B.-O., Wikby, A., Kipling, D., and Dunn-Walters, D.K. (2009). B-cell diversity decreases in old age and is correlated with poor health status. Aging Cell 8, 18–25.

Horns, F., Dekker, C.L., and Quake, S.R. (2020). Memory B Cell Activation, Broad Anti-influenza Antibodies, and Bystander Activation Revealed by Single-Cell Transcriptomics. Cell Rep. 30, 905–913.e6.

Joo, H.M., He, Y., and Sangster, M.Y. (2008). Broad dispersion and lung localization of virus-specific memory B cells induced by influenza pneumonia. Proceedings of the National Academy of Sciences 105, 3485–3490.

Koike, T., Harada, K., Horiuchi, S., and Kitamura, D. (2019). The quantity of CD40 signaling determines the differentiation of B cells into functionally distinct memory cell subsets. Elife 8.

Kometani, K., Nakagawa, R., Shinnakasu, R., Kaji, T., Rybouchkin, A., Moriyama, S., Furukawa, K., Koseki, H., Takemori, T., and Kurosaki, T. (2013). Repression of the transcription factor Bach2 contributes to predisposition of IgG1 memory B cells toward plasma cell differentiation. Immunity 39, 136–147.

Krishnamurty, A.T., Thouvenel, C.D., Portugal, S., Keitany, G.J., Kim, K.S., Holder, A., Crompton, P.D., Rawlings, D.J., and Pepper, M. (2016). Somatically Hypermutated Plasmodium-Specific IgM(+) Memory B Cells Are Rapid, Plastic, Early Responders upon Malaria Rechallenge. Immunity 45, 402–414.

Kuraoka, M., Schmidt, A.G., Nojima, T., Feng, F., Watanabe, A., Kitamura, D., Harrison, S.C., Kepler, T.B., and Kelsoe, G. (2016). Complex Antigens Drive Permissive Clonal Selection in Germinal Centers. Immunity 44, 542–552.

Kurosaki, T., Kometani, K., and Ise, W. (2015). Memory B cells. Nat. Rev. Immunol. 15, 149–159.

Le Gallou, S., Zhou, Z., Thai, L.-H., Fritzen, R., de Los Aires, A.V., Mégret, J., Yu, P., Kitamura, D., Bille, E., Tros, F., et al. (2018). A splenic IgM memory subset with antibacterial specificities is sustained from persistent mucosal responses. J. Exp. Med. 215, 2035–2053.

Lutz, J., Dittmann, K., Bösl, M.R., Winkler, T.H., Wienands, J., and Engels, N. (2015). Reactivation of IgG-switched memory B cells by BCR-intrinsic signal amplification promotes IgG antibody production. Nat. Commun. 6, 8575.

McCray, P.B., Jr, Pewe, L., Wohlford-Lenane, C., Hickey, M., Manzel, L., Shi, L., Netland, J., Jia, H.P., Halabi, C., Sigmund, C.D., et al. (2007). Lethal infection of K18-hACE2 mice infected with severe acute respiratory syndrome coronavirus. J. Virol. 81, 813–821.

McHeyzer-Williams, L.J., Milpied, P.J., Okitsu, S.L., and McHeyzer-Williams, M.G. (2015). Class-switched memory B cells remodel BCRs within secondary germinal centers. Nat. Immunol. 16, 296–305.

Mesin, L., Schiepers, A., Ersching, J., Barbulescu, A., Cavazzoni, C.B., Angelini, A., Okada, T., Kurosaki, T., and Victora, G.D. (2020). Restricted Clonality and Limited Germinal Center Reentry Characterize Memory B Cell Reactivation by Boosting. Cell 180, 92–106.e11.

Mimitou, E.P., Cheng, A., Montalbano, A., Hao, S., Stoeckius, M., Legut, M., Roush, T., Herrera, A., Papalexi, E., Ouyang, Z., et al. (2019). Multiplexed detection of proteins, transcriptomes, clonotypes and CRISPR perturbations in single cells. Nat. Methods 16, 409–412.

Moran, I., Nguyen, A., Khoo, W.H., Butt, D., Bourne, K., Young, C., Hermes, J.R., Biro, M., Gracie, G., Ma, C.S., et al. (2018). Memory B cells are reactivated in subcapsular proliferative foci of lymph nodes. Nat. Commun. 9, 3372.

Nakagawa, R., Toboso-Navasa, A., Schips, M., Young, G., Bhaw-Rosun, L., Llorian-Sopena, M., Chakravarty, P., Sesay, A.K., Kassiotis, G., Meyer-Hermann, M., et al. (2021). Permissive selection followed by affinity-based proliferation of GC light zone B cells dictates cell fate and ensures clonal breadth. Proc. Natl. Acad. Sci. U. S. A. 118.

Nojima, T., Haniuda, K., Moutai, T., Matsudaira, M., Mizokawa, S., Shiratori, I., Azuma, T., and Kitamura, D. (2011). In-vitro derived germinal centre B cells differentially generate memory B or plasma cells in vivo. Nat. Commun. 2, 465.

O’Flanagan, C.H., Campbell, K.R., Zhang, A.W., Kabeer, F., Lim, J.L.P., Biele, J., Eirew, P., Lai, D., McPherson, A., Kong, E., et al. (2019). Dissociation of solid tumor tissues with cold active protease for single-cell RNA-seq minimizes conserved collagenase-associated stress responses. Genome Biol. 20, 210.

Oh, J.E., Iijima, N., Song, E., Lu, P., Klein, J., Jiang, R., Kleinstein, S.H., and Iwasaki, A. (2019). Migrant memory B cells secrete luminal antibody in the vagina. Nature 571, 122–126.

Onodera, T., Takahashi, Y., Yokoi, Y., Ato, M., Kodama, Y., Hachimura, S., Kurosaki, T., and Kobayashi, K. (2012). Memory B cells in the lung participate in protective humoral immune responses to pulmonary influenza virus reinfection. Proceedings of the National Academy of Sciences 109, 2485–2490.

Pape, K.A., Taylor, J.J., Maul, R.W., Gearhart, P.J., and Jenkins, M.K. (2011). Different B cell populations mediate early and late memory during an endogenous immune response. Science 331, 1203–1207.

Riedel, R., Addo, R., Ferreira-Gomes, M., Heinz, G.A., Heinrich, F., Kummer, J., Greiff, V., Schulz, D., Klaeden, C., Cornelis, R., et al. (2020). Discrete populations of isotype-switched memory B lymphocytes are maintained in murine spleen and bone marrow. Nat. Commun. 11, 2570.

Siegrist, C.-A., and Aspinall, R. (2009). B-cell responses to vaccination at the extremes of age. Nat. Rev. Immunol. 9, 185–194.

Stoeckius, M., Zheng, S., Houck-Loomis, B., Hao, S., Yeung, B.Z., Mauck, W.M., 3rd, Smibert, P., and Satija, R. (2018). Cell Hashing with barcoded antibodies enables multiplexing and doublet detection for single cell genomics. Genome Biol. 19, 224.

Stone, S.L., Peel, J.N., Scharer, C.D., Risley, C.A., Chisolm, D.A., Schultz, M.D., Yu, B., Ballesteros-Tato, A., Wojciechowski, W., Mousseau, B., et al. (2019). T-bet Transcription Factor Promotes Antibody-Secreting Cell Differentiation by Limiting the Inflammatory Effects of IFN-γ on B Cells. Immunity 50, 1172–1187.e7.

Stuart, T., Butler, A., Hoffman, P., Hafemeister, C., Papalexi, E., Mauck, W.M., 3rd, Hao, Y., Stoeckius, M., Smibert, P., and Satija, R. (2019). Comprehensive Integration of Single-Cell Data. Cell 177, 1888–1902.e21.

Suan, D., Kräutler, N.J., Maag, J.L.V., Butt, D., Bourne, K., Hermes, J.R., Avery, D.T., Young, C., Statham, A., Elliott, M., et al. (2017). CCR6 Defines Memory B Cell Precursors in Mouse and Human Germinal Centers, Revealing Light-Zone Location and Predominant Low Antigen Affinity. Immunity 47, 1142–1153.e4.

Szabo, P.A., Miron, M., and Farber, D.L. (2019). Location, location, location: Tissue resident memory T cells in mice and humans. Sci Immunol 4.

Tamura, S.I., Asanuma, H., Ito, Y., Hirabayashi, Y., Suzuki, Y., Nagamine, T., Aizawa, C., Kurata, T., and Oya, A. (1992). Superior cross-protective effect of nasal vaccination to subcutaneous inoculation with influenza hemagglutinin vaccine. Eur. J. Immunol. 22, 477–481.

Trivedi, N., Weisel, F., Smita, S., Joachim, S., Kader, M., Radhakrishnan, A., Clouser, C., Rosenfeld, A.M., Chikina, M., Vigneault, F., et al. (2019). Liver Is a Generative Site for the B Cell Response to Ehrlichia muris. Immunity 51, 1088–1101.e5.

Viant, C., Weymar, G.H.J., Escolano, A., Chen, S., Hartweger, H., Cipolla, M., Gazumyan, A., and Nussenzweig, M.C. (2020). Antibody Affinity Shapes the Choice between Memory and Germinal Center B Cell Fates. Cell 183, 1298–1311.e11.

Viant, C., Wirthmiller, T., ElTanbouly, M.A., Chen, S.T., Cipolla, M., Ramos, V., Oliveira, T.Y., Stamatatos, L., and Nussenzweig, M.C. (2021). Germinal center-dependent and -independent memory B cells produced throughout the immune response. J. Exp. Med. 218.

Villazala-Merino, S., Rodriguez-Dominguez, A., Stanek, V., Campion, N.J., Gattinger, P., Hofer, G., Froeschl, R., Fae, I., Lupinek, C., Vrtala, S., et al. (2020). Allergen-specific IgE levels and the ability of IgE-allergen complexes to cross-link determine the extent of CD23-mediated T-cell activation. J. Allergy Clin. Immunol. 145, 958–967.e5.

Wan, Z., Lin, Y., Zhao, Y., and Qi, H. (2019). T cells in bystander and cognate interactions with B cells. Immunol. Rev. 288, 28–36.

Weisel, F., and Shlomchik, M. (2017). Memory B Cells of Mice and Humans. Annu. Rev. Immunol. 35, 255–284.

Weisel, F.J., Zuccarino-Catania, G.V., Chikina, M., and Shlomchik, M.J. (2016). A Temporal Switch in the Germinal Center Determines Differential Output of Memory B and Plasma Cells. Immunity 44, 116–130.

Wong, R., Belk, J.A., Govero, J., Uhrlaub, J.L., Reinartz, D., Zhao, H., Errico, J.M., D’Souza, L., Ripperger, T.J., Nikolich-Zugich, J., et al. (2020). Affinity-Restricted Memory B Cells Dominate Recall Responses to Heterologous Flaviviruses. Immunity 53, 1078–1094.e7.

Yeh, C.-H., Nojima, T., Kuraoka, M., and Kelsoe, G. (2018). Germinal center entry not selection of B cells is controlled by peptide-MHCII complex density. Nat. Commun. 9, 928.

Zaretsky, I., Atrakchi, O., Mazor, R.D., Stoler-Barak, L., Biram, A., Feigelson, S.W., Gitlin, A.D., Engelhardt, B., and Shulman, Z. (2017). ICAMs support B cell interactions with T follicular helper cells and promote clonal selection. Journal of Experimental Medicine 214, 3435–3448.

Zhou, P., Yang, X.-L., Wang, X.-G., Hu, B., Zhang, L., Zhang, W., Si, H.-R., Zhu, Y., Li, B., Huang, C.-L., et al. (2020). A pneumonia outbreak associated with a new coronavirus of probable bat origin. Nature 579, 270–273.

Zuccarino-Catania, G.V., Sadanand, S., Weisel, F.J., Tomayko, M.M., Meng, H., Kleinstein, S.H., Good-Jacobson, K.L., and Shlomchik, M.J. (2014). CD80 and PD-L2 define functionally distinct memory B cell subsets that are independent of antibody isotype. Nat. Immunol. 15, 631–637.

